# Phenotype shift scores reveal the scale and phylogenetic structure of phenotypic differentiation in primates

**DOI:** 10.64898/2026.09.03.749256

**Authors:** Fabio Barteri, Arcadi Navarro, Omar E. Cornejo

## Abstract

Continuous traits evolve unevenly across phylogenies, producing patterns of phenotypic differentiation shaped by both shared ancestry and lineage-specific change. Identifying exceptionally differentiated species pairs may therefore improve genome–phenome comparisons, but existing approaches rarely rank such contrasts across entire trees. Here we introduce the Phenotype Shift Score (PSS), a phylogenetic comparative framework integrating model-based trait divergence, observed trait range and patristic distance. Applying PSS to 153 continuous primate traits reveals trait-specific distributions of extreme differentiation across evolutionary depth. Body and brain mass show deeper-than-expected extremes under fitted Brownian motion or Ornstein–Uhlenbeck nulls, whereas body-size-adjusted brain mass localizes recent differentiation within cercopithecid lineages, complementing published branchwise reconstructions. PSS-informed groups produce more selective and more strongly enriched comparative-genomic signals than groups based on absolute phenotypic extremes. PSS is implemented in the open-source R package phyloPSS, with pairwise results available through the Primate Genome–Phenome Archive.

## 1. Introduction

The repeated occurrence of similar phenotypes in unrelated lineages is a pervasive pattern in the tree of life (Stayton 2015, Allard and Kumar 2026). Camera eyes in vertebrates and cephalopods, powered flight in birds, bats and insects, and body streamlining in aquatic tetrapods are the textbook cases, but convergence is at least as common at finer scales, in metabolism, diet, sensory capability, longevity and tolerance of environmental stress, where it is neither conspicuous nor easily catalogued (Losos 2011; Stern 2013). Beyond documenting repeatability and a degree of determinism in evolution (Blount et al. 2018), convergence has a practical value that has driven a decade of comparative genomics: when the same phenotype arises more than once in distantly related lineages, the genetic changes shared by those lineages, and absent from their close relatives, become candidates for the molecular basis of the trait (Stern 2013; Allard and Kumar 2026).

This logic underlies a large family of genome-wide scans and full research programs that include the search for what makes us human and what are the genetic bases of human specific phenotypes, including disease (Juan et al. 2023). Some search for identical or physicochemical similar amino acid substitutions at homologous sites in trait-bearing lineages (Zhang and Kumar 1997; Parker et al. 2013; Foote et al. 2015), an approach formalized for convergent amino acid substitutions in CAAStools (Barteri et al. 2024) and applied to traits such as mammalian life span (Muntané et al. 2018; Farré et al. 2021). Others look for shifts in evolutionary rate or in the strength of constraint that track the phenotype, as in forward genomics (Hiller et al. 2012), RERconverge (Chikina et al. 2016; Kowalczyk et al. 2019) and PhyloAcc (Hu et al. 2019). More recently, supervised learning has been used to fit sparse predictive models across entire proteomes, as in evolutionary sparse learning with paired species contrasts (Allard et al. 2025). These methods differ substantially in the molecular signal they target and in how they control for background convergence arising from shared history rather than adaptation (Castoe et al. 2009; Thomas and Hahn 2015; Zou and Zhang 2015). They share, however, a common dependency that has received far less attention than the molecular machinery built on top of it: they all require a priori definition of the phenotype shift, as well as the location in the phylogenetic tree for these shifts as starting points of the analysis.

That requirement is easy to satisfy for traits that are genuinely discrete, such as echolocation or C4 photosynthesis, and it is the reason those systems dominate the literature. It is much harder for the continuous traits that describe most of organismal biology, and the difficulty is especially acute in primates, where nearly every trait of interest is quantitative: body mass, endocranial volume, maximum life span, age at first reproduction, gestation length, weaning age, home range size, group size, sexual dimorphism and dietary quality. In practice such a trait is made usable by thresholding it, and the threshold is almost always chosen by hand, sometimes after regressing the trait on body mass and binning the residuals. Doing so discards the magnitude of the difference, makes the resulting gene lists sensitive to a decision with no principled justification, and, most importantly, ignores the phylogenetic structure and model of evolution of the trait itself. Two species may have nearly identical values because they diverged recently and have had little opportunity to differ, and two species may differ greatly simply because a great deal of time separates them. Whether an observed trait difference is large or small is not a property of the difference alone; it is only interpretable relative to what a model of trait evolution predicts for that particular pair of taxa given the tree (Felsenstein 1985). A partition built without that correction hands the downstream molecular scan a phenotype label that is partly a restatement of the phylogeny, which is precisely the confound those scans then work hard to remove.

Phylogenetic comparative methods supply the missing model. Brownian motion (Felsenstein 1985) and Ornstein-Uhlenbeck processes (Hansen 1997; Butler and King 2004; Beaulieu et al. 2012) provide explicit expectations for the covariance of trait values among tips, and a mature set of tools locates adaptive shifts on a tree under those models, including SURFACE (Ingram and Mahler 2013), bayou (Uyeda and Harmon 2014), l1ou (Khabbazian et al. 2016), PhylogeneticEM (Bastide et al. 2018) and the RRphylo shift and convergence searches (Castiglione et al. 2019, 2020). A parallel literature quantifies convergence itself, most prominently through Stayton’s C metrics (Stayton 2015) and the Wheatsheaf index (Arbuckle et al. 2014). These methods are well suited to the questions they were built for, but they are an awkward fit for genotype-phenotype mapping in two respects. First, their output is a shift configuration on branches, a regime assignment, or a single test statistic for a set of taxa designated in advance; none of these is the list of contrasting taxa that a molecular scan consumes, and converting between the two reintroduces the manual judgment we were trying to eliminate. Second, regime inference and shift-configuration search are computationally demanding and can be unstable, which becomes limiting when the goal is to screen many traits, large trees, non-ultrametric trees, or trait matrices with substantial missing data.

Here we introduce the Phenotype Shift Score (PSS), implemented in the open-source R package phyloPSS, a model-based index designed specifically to identify shifts in quantitative traits and facilitate molecular scans for the genetic changes underlying those shifts. For each trait we fit Brownian motion and single-optimum Ornstein-Uhlenbeck models and select between them by AIC under a conservative rule that retains Brownian motion unless the OU fit is clearly better, so that model support is never assumed where the data are ambiguous. From the fitted model we obtain, for every unordered pair of species, the expected variance of the tip contrast implied by the model covariance matrix, and we place the observed absolute difference on the resulting half-normal distribution. The result is a value between zero and one that states how extreme that pair’s phenotypic difference is relative to the amount of independent evolution separating the two taxa. This quantity is then combined with the difference normalized by the largest difference observed for the trait, and divided by the patristic distance normalized by the largest distance on the tree, so that pairs achieving a large phenotypic difference over a short evolutionary separation are ranked above pairs whose difference simply reflects deep divergence. PSS is a ranking index rather than a p-value, a probability of convergence, or an estimate of evolutionary rate, and we treat it throughout as a prioritization device rather than a hypothesis test.

Two design choices follow from the intended use. The score is defined on species pairs because the pair is the unit that downstream methods actually consume: CAAS discovery contrasts groups of species at homologous alignment positions (Barteri et al. 2024), and ESL-PSC is built on explicitly paired contrasts whose construction the authors identify as the principal practical obstacle to using the method (Allard et al. 2025). A ranked table of pairs can be passed directly into either framework. The score is also computed independently for each pair, so it parallelizes trivially and tolerates the missing data that is the norm in primate trait compilations, where a species may have a reliable body mass but no endocranial volume or no recorded maximum life span. Each trait is simply restricted to the taxa that have it, and no imputation or ultrametric assumption is required.

To showcase our method, we decided to use the Primate phylogeny, and associated rich phenotype datasets already explored in previous work (Kuderna et al. 2023, Galán-Acedo et al. 2019, Jones et al. 2009). We count with a high-coverage whole-genome data for 233 species spanning 86% of genera and all sixteen families, together with a fossil-calibrated nuclear phylogeny estimated from those same data (Kuderna et al. 2023), so the tree on which trait evolution is modeled and the alignments on which molecular convergence is scored come from a single, internally consistent resource. Comparative trait coverage is correspondingly deep, through curated primate-specific compilations (Galán-Acedo et al. 2019) and broader mammalian life-history and ageing databases (Jones et al. 2009; Tacutu et al. 2018). Primates also display exactly the kind of repeated quantitative transitions that motivate the method: independent expansions and reductions of relative brain size associated with dietary shifts (DeCasien et al. 2017), independent origins of extreme longevity relative to body size, and repeated transitions in activity pattern, locomotion and social organization across the Strepsirrhini, Platyrrhini and Catarrhini. Critically, primates are also the clade in which the genotype side of this pipeline has already been built out, with CAAS-based scans applied to life span and other life-history traits (Muntané et al. 2018; Farré et al. 2021; Barteri et al. 2024). Those studies had to define contrasting species groups by hand, which makes primates both the most natural demonstration of our approach and the case where a principled replacement for that step is most immediately useful. In what follows we describe the score and its behavior under simulation, characterizing how it responds to shifts of known magnitude and placement under both Brownian and OU generating processes, and how it compares with existing shift-detection and convergence measures. We apply PSS to the primate dataset and show that can recover well established phenotype shifts, as well as new ones(?), and we further show how the identification and proper coding of those shifts allow us to scan for convergent amino acid substitutions, yielding candidate gene changes explaining the evolution of the traits. Our aim is not to replace regime-based comparative methods but to provide the quantitative, phylogenetically explicit phenotype layer that genome-wide scans for convergence currently lack.

## 2. Results

### 2.1 Phenotype Shift Scores integrate model-calibrated extremeness, phenotypic magnitude and phylogenetic distance

We developed the Phenotype Shift Score (PSS) as a continuous pairwise statistic for ranking phenotypic differentiation across a phylogeny. For each trait, observed species were matched to the phylogeny, the tree was pruned to those species and every unordered pair was evaluated (Fig. 1a). The resulting comparison universe was used both to fit evolutionary expectations and to scale the observed quantities; species lacking measurements therefore contributed neither comparisons nor scaling maxima.

**Figure 1.**
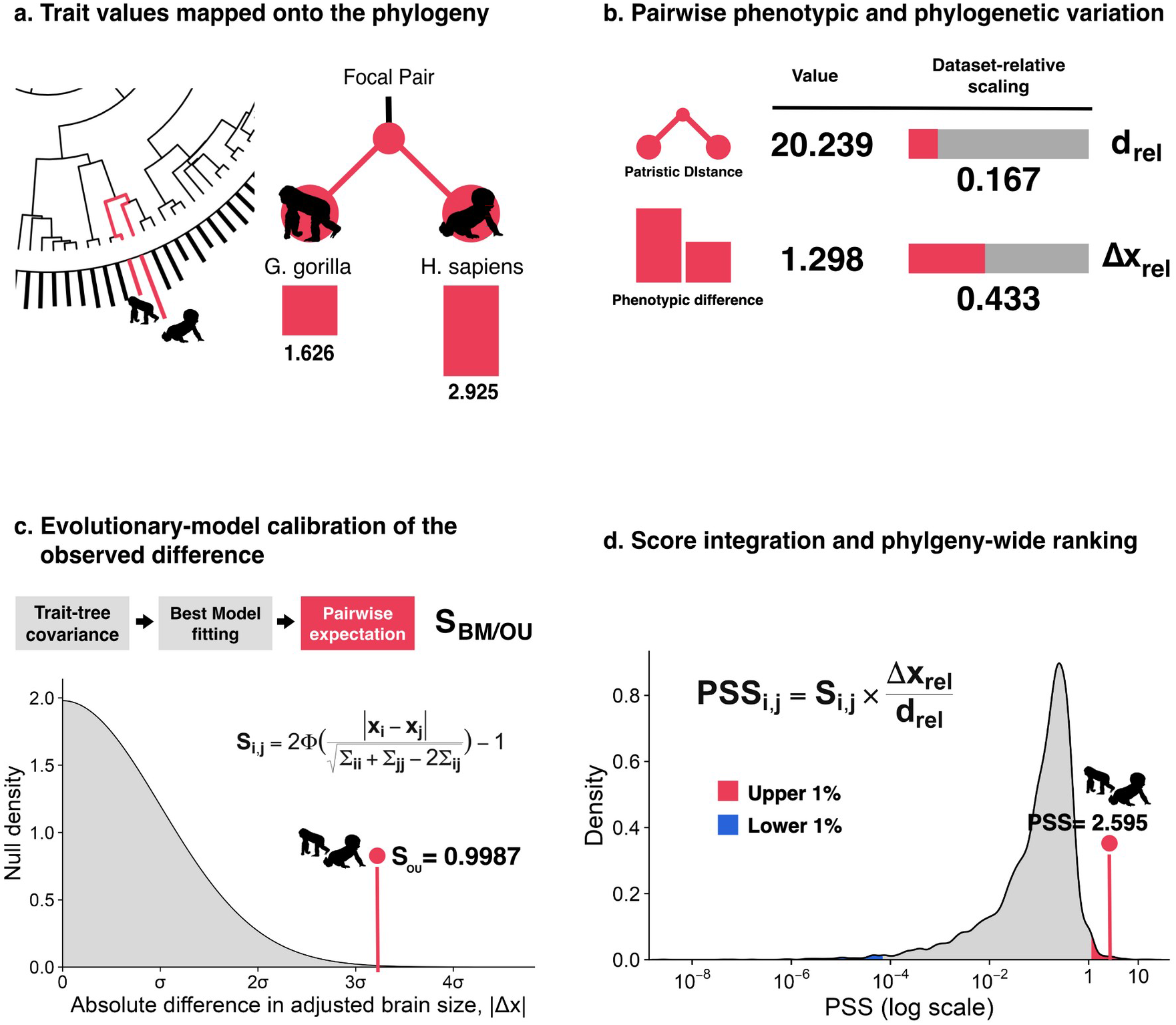
Model-informed calculation and phylogeny-wide ranking of phenotype shift scores. The phenotype shift score (PSS) ranks pairwise phenotypic contrasts by combining their magnitude, phylogenetic concentration and probability under a fitted evolutionary model. The illustrated calculation uses allometry-adjusted brain mass in 129 primates, with *Gorilla gorilla* and *Homo sapiens* as the focal pair. **a,** Trait values are mapped to the phylogeny before all unordered species pairs are evaluated. The focal species have trait values of 1.626 and 2.925, respectively. **b,** Pairwise phenotypic and phylogenetic variation. The focal pair has a patristic distance of 20.239 and an absolute phenotypic difference of 1.298. Each quantity is scaled by its maximum across the trait-specific dataset, yielding relative patristic distance *d*rel = 0.167 and relative phenotypic difference Δ*x*rel = 0.433. **c,** Evolutionary-model calibration of the observed difference. Brownian-motion and Ornstein–Uhlenbeck models are fitted to the trait, and the better-supported model determines the expected pairwise variance from the trait–tree covariance matrix. The absolute contrast is located in the corresponding half-normal distribution and transformed into *S*i,j, the model-calibrated extremeness of the pairwise difference; for the focal pair, the selected Ornstein–Uhlenbeck model gives *S*OU = 0.9987. **d,** Score integration and phylogeny-wide ranking. PSS is calculated as PSSi,j = *S*i,j(Δ*x*rel/*d*rel), favouring phenotypically extreme contrasts concentrated across comparatively short phylogenetic distances. The *Gorilla*–*Homo* comparison has PSS = 2.595 and ranks 22nd among 8,256 primate pairs, placing it in the Upper 1% of the trait-specific distribution. Upper and Lower 1% tails provide relative candidate sets for downstream analyses; PSS is a pairwise ranking index and does not identify the historical branch on which phenotypic change occurred.

The model-calibrated component, *S*, was derived from the joint distribution of tip values rather than from an empirical threshold. Under either a homogeneous Brownian-motion (BM) model or a homogeneous single-optimum Ornstein–Uhlenbeck (OU) model, the fitted trait vector is multivariate normal with a common expected tip value and a model-specific covariance matrix. Consequently, the signed contrast between species *i* and *j* is itself normally distributed with mean zero:

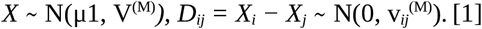

The variance expected for each contrast is pair-specific because it incorporates the two tip variances and their shared evolutionary history. If V^(M)^ is the covariance matrix implied by the fitted model, then

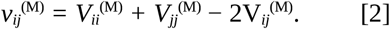

Under BM this reduces to the fitted evolutionary variance rate multiplied by the patristic distance, whereas under OU it follows the fitted attraction strength, diffusion variance and shared branch history and approaches a finite limit with increasing separation. BM and OU were fitted to the complete matched dataset. OU supplied the final covariance only when its AIC was more than two units lower than that of BM; otherwise the simpler BM model was retained (Fig. 1c).

Because the signed contrast has mean zero, its absolute value follows a half-normal distribution with scale √v*_ij_*^(M)^. We evaluated its cumulative distribution at the observed absolute difference δ*_ij_ = |x_i_ − x_j_*|. This yields the analytical placement score

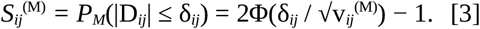

Thus, *S* is the percentile occupied by the observed difference in the theoretical absolute-difference distribution for that particular species pair. Values near one indicate that few differences predicted by the fitted model are as large as the observation, whereas values near zero indicate a difference that is small relative to the corresponding expectation. *S* is not a probability that divergence occurred and is not a pairwise hypothesis-test result; its analytical calculation avoids simulation and Monte Carlo error. The full derivation of S and its BM- and OU-specific variance is provided in Supplementary Note 2 (sections 2.1–2.2), with the analytical transformation and variance geometry shown in Supplementary Fig. 1a,b.

PSS then retained two descriptive quantities that *S* alone does not express. The observed absolute difference and patristic distance were each divided by their maximum across all available pairs, yielding dataset-relative phenotypic magnitude and relative phylogenetic distance (Fig. 1b):

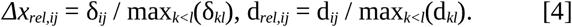

These dimensionless quantities preserve the ordering of the original measures while locating each comparison within the phenotypic and phylogenetic ranges sampled for that trait. The three components were integrated as

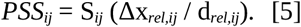

Accordingly, high values identify contrasts that are extreme under the selected evolutionary model, large relative to the observed phenotypic range and concentrated across comparatively short phylogenetic distances (Fig. 1d). In the worked primate example, *Gorilla gorilla* and *Homo sapiens* had allometry-adjusted brain-mass values of 1.626 and 2.925, respectively. Their absolute difference of 1.298 across a patristic distance of 20.239 corresponded to Δx*_rel_* = 0.433 and d*_rel_* = 0.167. OU was selected and gave S*_ij_* = 0.9987; the integrated PSS of 2.595 ranked 22nd among 8,256 pairs, placing the comparison in the Upper 1%. The full worked calculation, exact computational audit and matched pair controls are reported in Supplementary Note 2 (section 2.3), the accompanying Supplementary Table 1 and Supplementary Fig. 1d.

PSS therefore provides a continuous ranking rather than a binary test. The Upper and Lower 1% are operational candidate sets, not statistically significant classes, and an individual score is neither a p-value nor an estimate of evolutionary rate. Nor does PSS reconstruct ancestral states or identify the historical branch on which change occurred. Because phylogenetic distance enters first through the variance used to calculate *S* and again as an explicit denominator, the score intentionally gives additional weight to differentiation across short relative distances. The phylogenetic distribution of its tails must therefore be interpreted against trait-specific opportunity and fitted-model expectations. Component geometry, component-ablation results and tail-width sensitivity are detailed in Supplementary Note 2 (sections 2.4–2.5) and Supplementary Fig. 1c,e,f.

### 2.2 Extreme phenotypic differentiation occupies trait-specific phylogenetic depths in primates

We therefore evaluated PSS across 153 continuous traits retained from the Primate Genotype–Phenotype Archive (PGA; Valenzuela et al., 2026). Species coverage differed among traits, so each analysis used a separately pruned phylogeny, independently fitted and selected BM or OU model, and all pairwise comparisons available for that trait (Supplementary Note 3, sections 3.1–3.2; Supplementary Fig. 2). The Upper and Lower 1% of each score distribution again defined candidate sets rather than intrinsically significant classes.

We first expressed the median phylogenetic depth of each tail relative to all pair opportunities on the corresponding pruned tree (Fig. 2a), with 0% denoting the shallowest available comparisons, 50% the median of all available pairs and 100% the deepest. Across traits, the median of the trait-specific Upper 1% depth medians was 2.96% (interquartile range, 1.59–13.33%), compared with 55.93% (39.91–63.33%) for the Lower 1%. Upper-tail pairs were thus generally concentrated among relatively shallow comparisons. This tendency is expected from the inverse distance term in PSS and cannot by itself be interpreted as a universal biological preference for recent differentiation.

**Figure 2.**
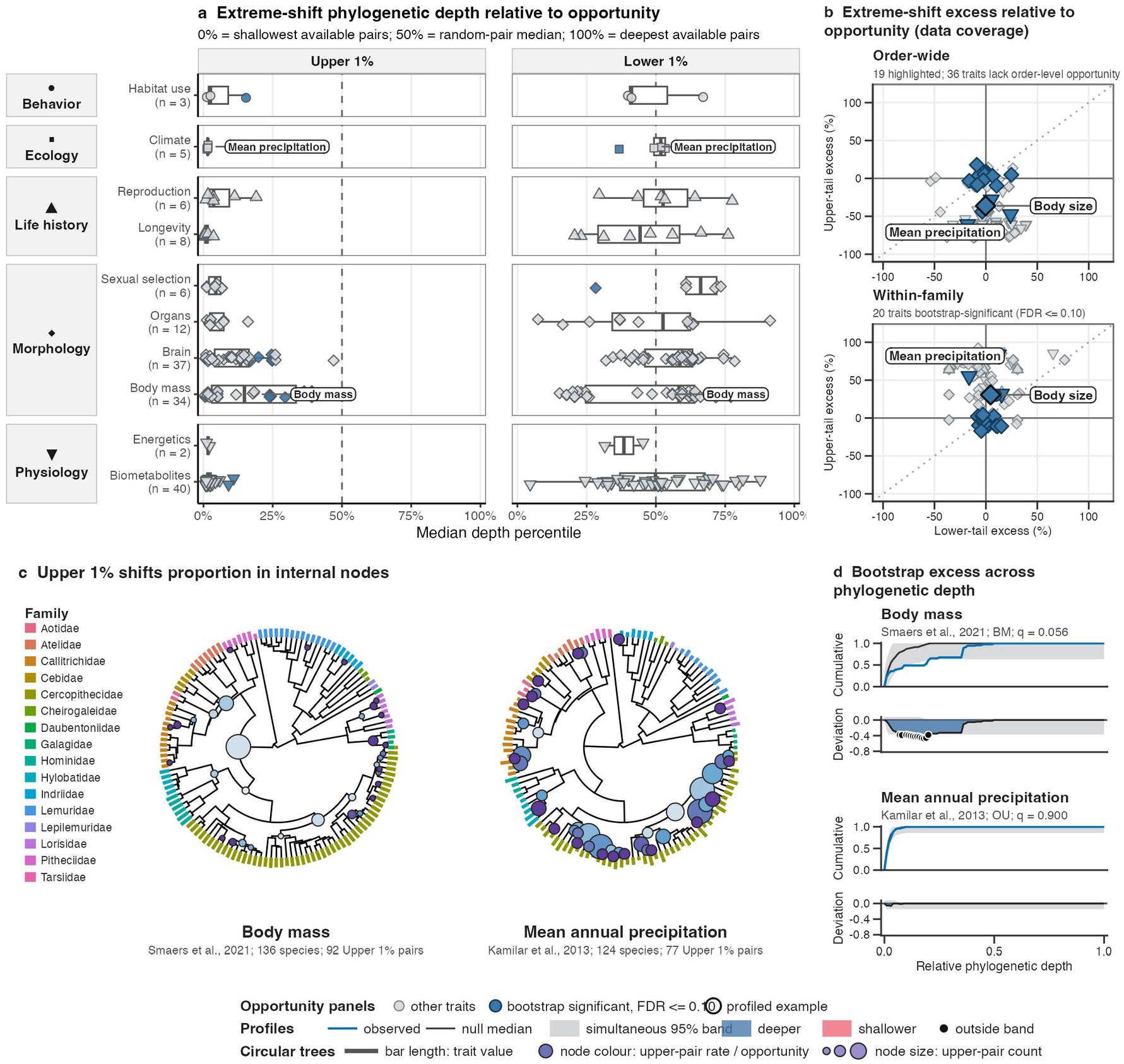
Phylogenetic distribution of extreme phenotype shifts in primate traits. Phenotype shift scores (PSSs) were calculated for 153 traits retained from the Primate Genotype–Phenotype Archive (PGA; Valenzuela et al., 2026). Extreme shifts were species pairs in the Upper 1% or Lower 1% of each trait’s PSS distribution. **a,** Tail phylogenetic depth relative to trait-specific pairwise opportunity. Percentiles of 0%, 50% and 100% denote the shallowest pairs, random-pair median and deepest pairs, respectively. Points represent traits, boxplots summarize PGA subdomains and symbols identify domains. Blue marks tail-specific profiles significant in the full bootstrap at false-discovery rate (FDR) 0.10. **b,** Opportunity-adjusted coverage at the order-wide and within-family levels. Excess equals the tail share at the focal level minus its share among all available pairs; zero denotes proportional representation. Axes give Lower 1% and Upper 1% excess. Dark-blue points have a significant negative signed area in the Upper 1% bootstrap depth profile, indicating redistribution towards deeper comparisons; labelled traits are examined in **c** and **d**. **c,** Nodal localization of Upper 1% shifts for body mass (Smaers et al., 2021) and mean annual precipitation (Kamilar et al., 2013). Each pair was assigned to its most recent common ancestor. Node size gives the number of assigned shifts and colour their proportion among nodal pairwise opportunities. External bars encode relative trait value by length and family by colour. Body-mass shifts accumulate at deeper nodes joining distant species, whereas precipitation shifts concentrate among shallower nodes joining close relatives. **d,** Full parametric-bootstrap depth profiles. Evolution was simulated under the fitted Brownian-motion (BM) or Ornstein–Uhlenbeck (OU) model, after which models were refitted and reselected and PSSs recalculated. Upper plots compare observed cumulative depth with the bootstrap median and simultaneous 95% envelope; lower plots give observed-minus-null deviation, with negative values indicating deeper redistribution. Body mass departs towards deep Upper 1% shifts (BM; FDR-adjusted *q* = 0.056), whereas mean annual precipitation follows its fitted OU expectation (*q* = 0.900).

Trait coverage also changed which taxonomic comparisons were possible. To separate observed tail composition from this opportunity, we calculated order-wide and within-family excess as the fraction of tail pairs at the focal level minus its fraction among all tested pairs for the same trait; zero therefore indicates proportional representation (Fig. 2b). Thirty-six traits lacked order-level opportunity and were not estimable at that scale (Supplementary Note 3, section 3.3; Supplementary Fig. 3). Excess varied widely within PGA domains and subdomains, and no primary domain was uniformly associated with a single phylogenetic depth. For example, the Body mass Upper 1% was enriched within families by 30.6 percentage points and depleted order-wide by 36.2 points, whereas mean annual precipitation showed corresponding excesses of +82.0 and −70.9 points. These opportunity-adjusted summaries remained descriptive and did not establish departure from the fitted evolutionary process.

We tested for such departures using 1,000 full parametric-bootstrap replicates per trait. Each replicate simulated phenotypes under the selected BM or OU model, refitted and reselected both candidate models, recalculated every PSS value and reapplied the same 1% tail definition. Observed cumulative depth profiles were compared with the bootstrap median and simultaneous 95% envelope, and empirical probabilities for absolute profile area were corrected across traits. At a false-discovery rate (FDR) of 0.10, 20 Upper 1% profiles combined a negative signed area with a significant absolute-area departure, indicating a redistribution towards deeper comparisons relative to the fitted null. All 20 were estimable within families and 19 at the order-wide level. They comprised 11 body-mass measurements, five brain measurements, three biometabolite traits and territoriality; repeated body- and brain-mass definitions provide robustness across related measurements rather than independent biological replication. Two Lower 1% profiles—mid-range latitude and male canine height—showed the converse, shallower-than-null redistribution (Supplementary Note 3, sections 3.4–3.5; Supplementary Fig. 4).

Body mass and mean annual precipitation illustrated why raw taxonomic or nodal localization required this model calibration (Fig. 2c,d). Body mass (Smaers et al., 2021) included 136 species, 9,180 pairs and 92 Upper 1% pairs, with BM selected. Its observed cumulative profile fell below the refitted-null profile over restricted depth intervals, yielding a signed area of −0.126 and an FDR-adjusted q = 0.056. In contrast, mean annual precipitation (Kamilar and Cooper, 2013) included 124 species, 7,626 pairs and 77 Upper 1% pairs. OU was selected in every bootstrap replicate, the signed area was −0.0019 (q = 0.900), and the observed profile never crossed the simultaneous envelope. Thus, body mass showed a deepward departure from its fitted BM null, whereas the visually shallower precipitation pattern was consistent with its fitted OU expectation.

The accompanying MRCA maps localized the extant comparisons contributing to these profiles without assigning historical change to particular branches. Node size represented the number of Upper 1% pairs assigned once to that node, whereas node colour expressed this count relative to nodal pair opportunity. Because the bootstrap retained depth distributions rather than simulated node identities, these visualizations provide descriptive localization rather than node-level significance. The phenome-wide calibration therefore identified traits whose extreme differentiation departed from score geometry and model-based expectation, but it did not yet show whether PSS could separate correlated phenotypes and preserve structure across nested taxonomic scales.

### 2.3 Brain–body allometry reveals multiscale structure in extreme phenotypic differentiation

We addressed this question using log body mass, log brain mass and allometry-adjusted brain mass for 1,427 mammals from Venditti et al. (2024). Each trait generated 1,017,451 unordered pairs and 10,175 Upper 1% pairs. Their taxonomic composition differed markedly among traits and orders (Fig. 3a). Among pairs involving Primates, 416 of 431 body-mass pairs (96.5%) and 519 of 521 brain-mass pairs (99.6%) were within-order comparisons, compared with 1,057 of 1,576 pairs (67.1%) for adjusted brain mass. Variation was also evident across Carnivora, Rodentia and Cetacea, demonstrating that the upper-tail distributions were not taxonomically interchangeable. These fractions describe tail composition and do not directly imply accelerated evolution (Supplementary Note 4, sections 4.1–4.2; Supplementary Fig. 5; Source Data 2).

**Figure 3.**
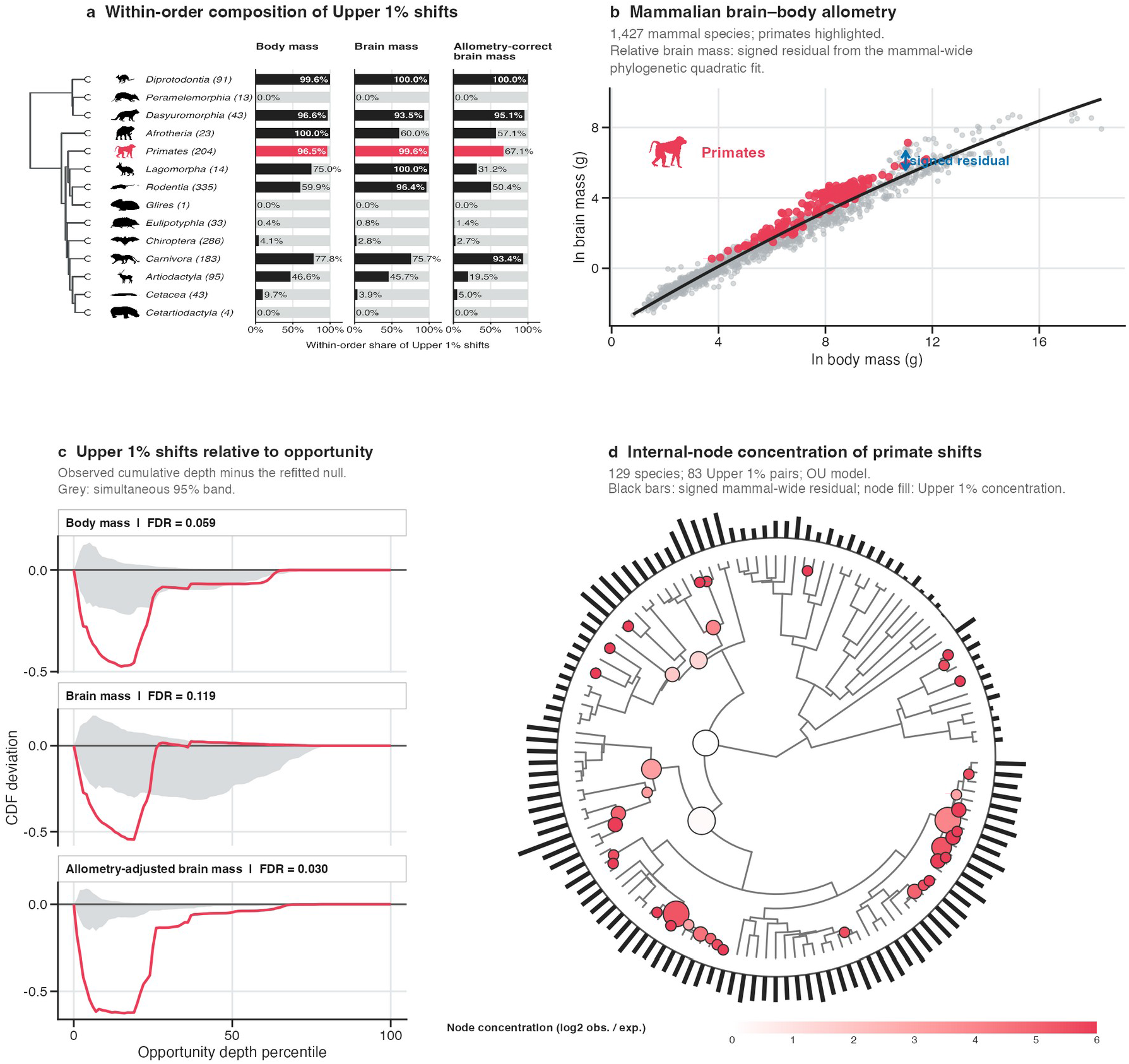
Mammalian distribution and primate localization of extreme brain–body phenotype shifts. Phenotype shift scores (PSSs) were compared for log body mass, log brain mass and allometry-adjusted brain mass across 1,427 mammals (Venditti et al., 2024). Extreme shifts are species pairs in the Upper 1% of each trait-specific PSS distribution. **a,** Within-order representation of Upper 1% shifts for the three traits. For each order, the grey bar denotes all Upper 1% pairs involving at least one member of that order (normalized to 100%), whereas the superimposed black bar gives the percentage for which both species belong to the order. Primates and their within-order fractions are highlighted in red; numbers in parentheses give species coverage. Orders without Upper 1% representation in one or more traits are omitted. **b,** Mammalian brain–body allometry. Points show log brain mass against log body mass, the black curve is the mammal-wide phylogenetic quadratic fit and red points identify primates. Allometry-adjusted brain mass is the signed residual from this relationship. **c,** Full parametric-bootstrap validation of Upper 1% phylogenetic depth. Red curves give the observed cumulative-depth deviation from the refitted null, and grey areas are simultaneous 95% null envelopes. Negative deviations indicate a redistribution towards deeper comparisons after accounting for pairwise opportunity. False-discovery-rate-adjusted values are 0.059 for body mass, 0.119 for brain mass and 0.030 for allometry-adjusted brain mass. **d,** Primate-specific refit and internal-node localization of relative-brain-mass shifts. The mammal-wide signed residual was retained as the trait, after which the evolutionary model was refitted and PSSs recalculated across 129 primates; an Ornstein–Uhlenbeck model was selected and 83 pairs formed the Upper 1%. External black bars encode the signed residual, with outward and inward extension indicating positive and negative values, respectively. Each Upper 1% pair is assigned to its most recent common ancestor. Node size gives the number of assigned pairs, and fill gives their log2 concentration relative to nodal pairwise opportunity.

To distinguish relative brain mass from body-size variation, we fitted a mammal-wide phylogenetic quadratic relationship between log brain mass and log body mass (Fig. 3b). The signed residual defined allometry-adjusted brain mass: positive values indicated brain masses above the mammal-wide expectation at a given body mass, and negative values indicated values below it. This residual was derived once across all mammals and transferred unchanged to the nested primate analysis, preserving a common allometric reference rather than redefining the phenotype within each clade. We retained the quadratic specification as the reference phenotype because it follows the curvilinear brain–body relationship supported by Venditti et al. (2024). Diagnostic assessment nevertheless favoured a cubic relationship, although quadratic and cubic residual ranks remained highly concordant; exact tail rankings therefore remain conditional on model form (Supplementary Note 4, section 4.3; Supplementary Fig. 6).

Full parametric-bootstrap validation further separated the three phenotypes after accounting for pairwise opportunity and refitting and reselecting the evolutionary model in every replicate (Fig. 3c). Negative observed-minus-null cumulative-depth deviations indicated deepward redistribution. Log body mass had a signed area of −0.118 and *q* = 0.059; log brain mass had a smaller deviation (−0.094, *q* = 0.119); and adjusted brain mass showed the strongest departure (−0.159, *q* = 0.030). At FDR ≤ 0.10, body mass and adjusted brain mass therefore departed from their fitted null depth profiles, whereas raw brain mass did not. Because the residual is derived from brain and body mass, these are not independent trait tests; their contrast instead indicates that removing mammal-wide body-size allometry exposed phylogenetic-depth structure that was not statistically supported for raw brain mass alone.

We then retained the unchanged mammal-wide residual, pruned the Kuderna et al. (2023) phylogeny to 129 primates with matched observations and refitted BM and OU on this primate-specific tree (Fig. 3d). OU was selected, and recalculation across 8,256 available pairs identified 83 Upper 1% comparisons. Assigning each pair once to its MRCA revealed a non-uniform internal-node distribution: node size summarized assigned-pair counts, while fill represented their log2 concentration relative to nodal pair opportunity. This mapping localizes the separations across which extreme extant differences are concentrated; it does not identify a historical shift branch or supply node-specific bootstrap significance (Supplementary Note 4, section 4.4; Supplementary Fig. 7a; Source Data 2).

Much of the within-family upper-tail structure localized to Cercopithecidae, with 43 observed pairs compared with 14.929 expected from the family’s share of pairwise opportunity. Other represented families each contributed between one and five within-family pairs. Although this is a descriptive opportunity comparison rather than a family-specific bootstrap test, it defined a focused domain for downstream analysis. In a separate within-Cercopithecidae refit (Supplementary Note 4, sections 4.5–4.6; Supplementary Fig. 7b–f; Source Data 2), 55 species produced 1,485 pairs; OU was again selected, and 14 of the 15 Upper 1% pairs were congeneric, including comparisons within *Macaca*, *Papio* and *Trachypithecus*. The mammalian calibration and nested primate localization thus translated a class-wide allometric signal into phylogenetically explicit candidate comparisons for genome–phenome analysis.

### 2.4 PSS-informed comparisons generate more selective genome–phenome hypotheses

The concentration of high-ranking relative-brain-mass contrasts within Cercopithecidae provided the basis for four convergent amino acid substitution (CAAS) discovery designs on the same 129-species phylogeny and phenotype scale (Fig. 4a): primate-wide absolute phenotypic extremes, family-level parallel phenotypic pairs, Cercopithecidae foreground (FG) and background (BG) pools defined from high-ranking PSS contrasts, and Cercopithecidae-wide absolute phenotypic extremes. The last design was the closest control for PSS-informed selection because it held the focal family constant while changing the group-selection criterion. Each strategy was assessed through 100 unique pooled CAAStools cycles (Barteri et al., 2023), sampling four FG and four BG species per cycle. These cycles were alternative realizations from finite pools, not statistically independent experimental replicates.

**Figure 4.**
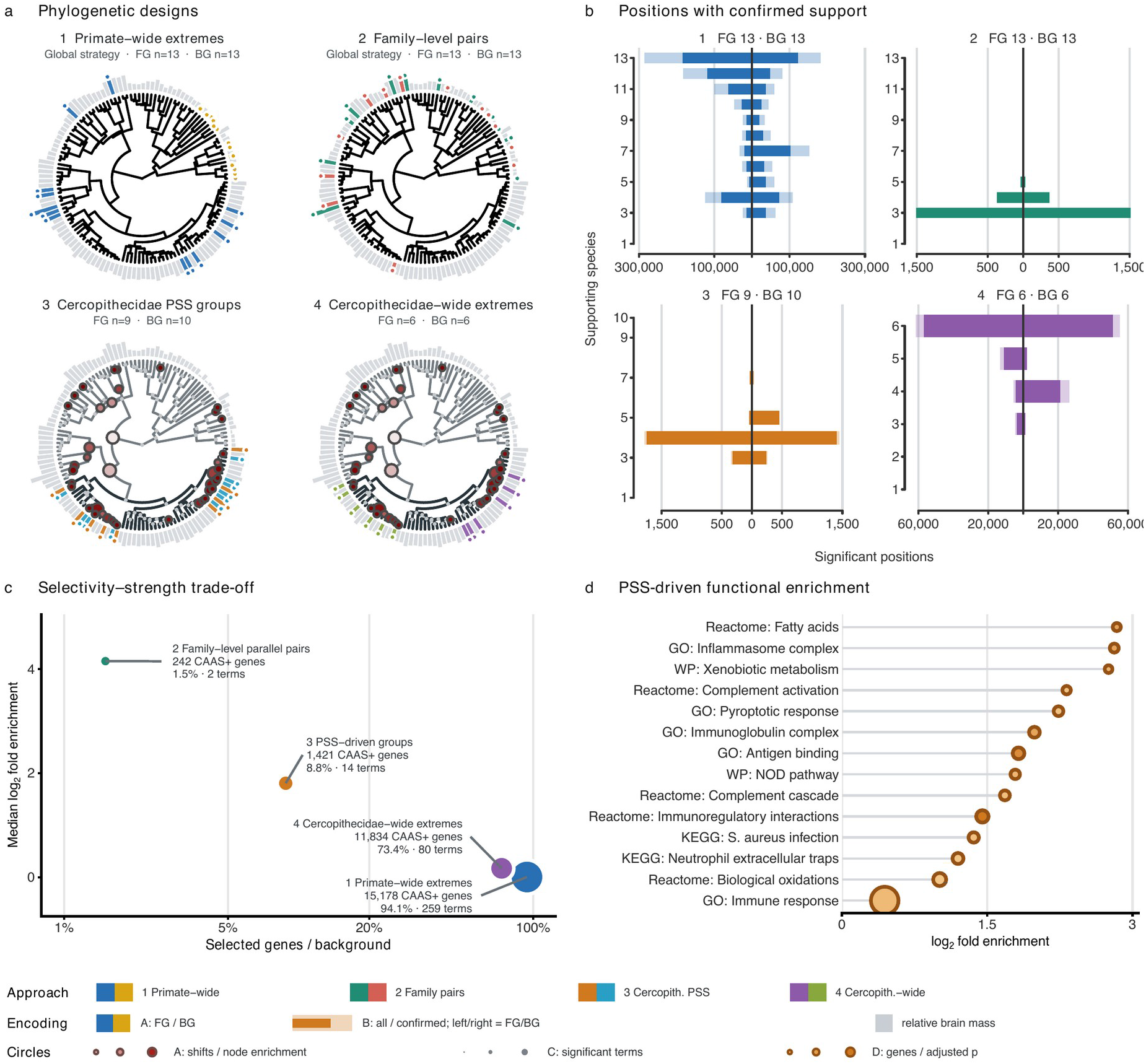
PSS-informed discovery-group design for CAAS detection and functional enrichment in relative-brain-mass evolution a, Foreground (FG) and background (BG) species pools for four convergent amino acid substitution (CAAS) discovery-group designs on the same 129-species primate phylogeny and relative-brain-mass scale: (1) primate-wide phenotypic extremes, (2) family-level parallel phenotypic pairs, (3) Cercopithecidae groups defined by high-ranking phenotype shift score (PSS) contrasts and (4) Cercopithecidae-wide phenotypic extremes. Grey radial bars denote unselected species; coloured bars and terminal circles identify FG and BG pools. Internal-circle size gives the number of global Upper 1% PSS shifts assigned to each node and fill gives log2 nodal enrichment relative to pairwise opportunity. **b,** Mirrored distributions of nominally significant CAAS positions (*P* < 0.05) by the number of supporting species. FG and BG support are shown to the left and right, respectively; pale bars show all positions and dark bars those retained after gene-wise dense-cluster filtering. Each strategy comprises 100 unique pooled CAAStools cycles, with four FG and four BG species sampled per cycle. **c,** Candidate-set selectivity and functional-enrichment strength. The horizontal axis gives the percentage of the strategy-specific tested background retained in the cluster-filtered, balanced-support query set; the vertical axis gives median log2 fold enrichment across significant g:Profiler terms. Circle area denotes the number of significant terms; labels report the number of CAAS+ query genes, query percentage and term count. **d,** Fourteen significant g:Profiler terms recovered by the PSS-driven strategy, ordered by log2 fold enrichment. Stems originate at no enrichment, circle area denotes intersecting query genes and colour denotes −log10 of the g:SCS-adjusted *P* value. In **c** and **d**, fold enrichment uses the effective strategy-specific custom background and includes only terms passing g:SCS-adjusted *P* < 0.05.

Primary CAAS discoveries were initially retained at nominal positional *P* < 0.05 and summarized by FG and BG species support (Fig. 4b). Primate-wide and Cercopithecidae-wide absolute extremes generated substantially broader positional sets than the family-pair and PSS-informed designs. We then removed dense gene-wise clusters of adjacent discoveries, which can arise from one alignment irregularity, indel boundary or extended sequence feature. The filter retained 63.38% of positions from primate-wide extremes and 89.35% from Cercopithecidae-wide extremes, compared with 99.64% from family-level pairs and 97.14% from PSS-informed groups. The PSS-informed signal was therefore predominantly composed of isolated positions and was minimally affected by this quality-control step. Because high discovery volume can also reflect genomic saturation, raw position or gene counts were not interpreted as evidence of superior performance.

For functional analysis, genes were retained only when a cluster-filtered significant position was supported at the same residue by at least four FG and four BG species. This produced 15,178 query genes for primate-wide extremes, 242 for family-level pairs, 1,421 for PSS-informed groups and 11,834 for Cercopithecidae-wide extremes, equivalent to 94.1%, 1.5%, 8.8% and 73.4% of the corresponding tested backgrounds (Fig. 4c). Using strategy-specific custom backgrounds, g:Profiler (Kolberg et al., 2023) recovered 259, 2, 14 and 80 terms at g:SCS-adjusted *P* < 0.05, with median fold enrichments of approximately 1.00, 18.59, 3.50 and 1.13, respectively. Absolute-tail designs therefore recovered many terms but selected most of their available backgrounds and produced effects near unity. Family-level pairs were exceptionally selective and strongly enriched but recovered only two functions. The PSS-informed design occupied an intermediate region, combining a restricted candidate set with appreciable functional breadth and stronger enrichment than either absolute-tail design.

The direct comparison within Cercopithecidae isolated the practical contribution of PSS-informed selection. Relative to absolute phenotypic extremes in the same family, PSS reduced the query from 73.4% to 8.8% of the tested background while increasing median enrichment from approximately 1.13-fold to 3.50-fold. This improvement coincided with a reduction from 80 to 14 significant terms rather than an increase in annotation count. PSS therefore did not maximize any single metric; it yielded a more favourable balance among candidate-set selectivity, enrichment strength and functional breadth.

The 14 PSS-associated terms included fatty-acid metabolism, inflammasome and pyroptotic responses, complement activation and cascade regulation, immunoglobulin complexes and antigen binding, NOD signalling, neutrophil extracellular traps, biological oxidation and xenobiotic metabolism (Fig. 4d). These enrichments provide structured and testable hypotheses spanning immune, inflammatory and metabolic systems, but they do not establish that particular substitutions caused relative-brain evolution. Together, the four analyses show that PSS can move from phylogeny-wide phenotypic ranking and model calibration to the construction of selective, biologically interpretable genome–phenome comparisons.

## 3. Discussion

The evolutionary interpretation of phenotypic differentiation depends on both the phylogenetic distance separating taxa and the trait-specific process generating variation. PSS addresses this problem by integrating phenotypic magnitude, phylogenetic separation and model-based extremeness on a common scale (Fig. 1; Supplementary Notes 1–2 and Supplementary Figure 1). Across primate traits, extreme comparisons occupied trait-specific phylogenetic depths, and several empirical profiles departed from fitted expectations after pairwise opportunity was accounted for (Fig. 2; Supplementary Note 3 and Supplementary Figures 2–4). The brain–body case study then showed how phenotype definition and fitting scale can reorganize this structure, whereas the Cercopithecidae benchmark demonstrated its practical consequence for genome–phenome hypothesis formulation (Figs. 3–4; Supplementary Notes 4–5 and Supplementary Figures 5–9). These results support a view of PSS as a discovery layer: it identifies where exceptional terminal contrasts are concentrated and which species configurations merit more focused evolutionary or genomic analysis.

PSS occupies a distinct but complementary position relative to established phylogenetic comparative approaches. Phylogenetic independent contrasts standardize nodal contrasts for association analyses (Felsenstein, 1985; Garland et al., 1992), historical convergence measures reconstruct whether focal lineages became more similar through time (Stayton, 2015; Grossnickle et al., 2024), and methods such as SURFACE, l1ou, bayou and phyloEM infer shifts among branches, adaptive regimes or evolutionary optima (Ingram and Mahler, 2013; Uyeda and Harmon, 2014; Khabbazian et al., 2016; Bastide et al., 2018). PSS addresses a different question. Among taxa retained after trait filtering and phylogenetic reconciliation, it evaluates all terminal-pair comparisons without requiring candidate species groups or specific regime shifts to be defined in advance, while still conditioning phenotypic differentiation on a fitted background model—either Brownian motion or an Ornstein–Uhlenbeck process (Supplementary Note 1 and Supplementary Table 1). The phenome-wide analysis illustrates the value of this distinction. Body-size and morphological traits tended to contribute contrasts spanning deeper divergences, whereas several ecological and physiological traits concentrated their most extreme comparisons within families or genera. Because these differences persisted after opportunity correction and full model refitting in the parametric bootstrap, they cannot be attributed solely to uneven taxon sampling or the greater availability of comparisons at particular depths. They instead indicate that different components of primate phenotypic diversity remain informative at different phylogenetic scales. Depth is not chronology, however: a concentration at deeper comparisons does not identify when a trait changed, locate a causal branch or imply an evolutionary-rate shift.

The brain–body case study further shows that phenotype construction is part of the biological hypothesis rather than a neutral preprocessing choice. Raw body mass, raw brain mass and mammal-wide allometry-adjusted brain mass produced different taxonomic and phylogenetic-depth profiles, although they were derived from closely related measurements (Fig. 3; Supplementary Note 4 and Supplementary Figure 5). Only the adjusted phenotype showed an FDR-supported redistribution towards deeper comparisons after full bootstrap calibration. Allometric adjustment therefore revealed structure that was not evident from raw brain mass, but it should not be interpreted as producing a uniquely correct measure of brain evolution. Empirical-tail membership remained partly sensitive to the allometric model, even though the signal was not driven by *Homo sapiens* or Hominidae (Supplementary Figure 6). Holding the mammal-wide phenotype definition constant while refitting the evolutionary model first to primates and then to Cercopithecidae localized the signal across nested phylogenetic scales and ultimately concentrated it in congeneric comparisons (Supplementary Figure 7 and Source Data 2). The case study therefore demonstrates both the value and the responsibility of phenotype definition: PSS can reveal structure hidden by raw measurements, but the resulting comparisons remain conditional on how the continuous trait is constructed.

The genome–phenome benchmark was motivated by a phenotypic configuration that absolute trait rankings would not have revealed. The primate-wide analysis localized a disproportionate concentration of high-ranking relative brain-mass comparisons within Cercopithecidae, and the family-specific refit showed that nearly all Upper 1% pairs were congeneric (Fig. 3; Supplementary Note 4 and Source Data 2). These species were not selected because they simply occupied opposite ends of the phenotype distribution. PSS instead identified a structured network of closely related taxa participating in unusually differentiated and unusually similar comparisons under the fitted model. A top-versus-bottom design would therefore have missed the combination of family-level concentration, congeneric differentiation and pairwise context that motivated the genomic analysis. Importantly, this structure was identified before the sequence results were examined and was used to formulate the foreground and background pools. Under the same pooled CAAStools workflow, the PSS-informed strategy selected a substantially smaller fraction of the tested gene background than absolute phenotype tails, reducing genomic saturation while retaining a non-trivial and functionally structured enrichment signal (Fig. 4; Supplementary Note 5 and Supplementary Figures 8–9). A smaller gene set is not intrinsically better, and the two strategies did not recover wholly separate gene catalogues. The relevant gain was instead conceptual: information extracted from the phenotypic network made it possible to formulate a more selective and discriminating genome–phenome hypothesis before sequence testing.

The interpretive advantages of PSS should be considered together with the assumptions and limitations of each analytical stage. Empirical PSS tails are prioritization sets rather than collections of independently significant species pairs, because taxa recur across comparisons and pairwise opportunity varies with phylogenetic depth. Statistical support therefore applies to aggregate profiles evaluated through the refitted parametric bootstrap, not automatically to individual pairs or clades (Fig. 2; Supplementary Note 3 and Supplementary Figures 3–4). A further consideration arises from the composite structure of the score. An extreme PSS does not require its model-based placement *S*, absolute phenotypic difference *Δ* or phylogenetic distance *T* to be individually extreme; it can result from the joint contribution of moderately large values across components (Fig. 1; Supplementary Note 2 and Supplementary Figure 1). Depending on the biological question, users may therefore supplement PSS-tail selection with explicit component-level criteria, such as a minimum phenotypic difference, a specified range of phylogenetic distances or a more extreme model-based placement. Because *S*, *Δ* and *T* are retained separately, such filtering can be applied transparently without redefining PSS. Rankings nevertheless remain conditional on trait coverage, taxonomic reconciliation and evolutionary-model choice. The brain–body analysis showed that alternative allometric models can affect membership near empirical-tail thresholds (Supplementary Figure 6). The genomic benchmark additionally depends on alignment quality, species coverage, cluster filtering and balanced-support requirements; pooled cycles are alternative realizations of finite species pools rather than independent evolutionary replicates (Supplementary Note 5, Section 5.5; Supplementary Figures 8–9). Finally, enriched immune and metabolic categories remain hypotheses: neither PSS nor pooled CAAStools identifies causal substitutions, demonstrates adaptation or establishes a mechanistic relationship between sequence and phenotype.

Taken together, PSS provides a scale-aware framework for converting continuous-trait variation into explicit comparative hypotheses. Its contribution is not a new reconstruction of evolutionary history, but an exhaustive, model-calibrated and bidirectional map of terminal differentiation that preserves species identities and exposes the phylogenetic scale of exceptional comparisons. The primate analyses show that this map can reveal trait-specific structure, localize non-obvious species networks and narrow the design space for downstream genome–phenome analyses. PSS therefore offers a transparent bridge between comparative phenomics and focused tests of evolutionary process and molecular association.

4.Methods

### Note on data and code availability

All software and resources supporting this study are openly available through the phyloPSS GitHub repository. The R package provides the reference implementation of PSS, while analysis pipelines, configurations, processed datasets and results are organized in the supplementary/ directory of the same repository. The archived release and persistent DOI will be added after deposition: [ADD ARCHIVE/DOI].

### 4.1 Phenotypic data and phylogeny

#### 4.1.1 Primate trait dataset

The broad phenome analysis used the Primate Genome Atlas (PGA) phenomic dataset stored in the project as nhp.phenomic.dataset.tsv (Kuderna et al., 2023). Species were identified by the SpeciesBROAD field. Trait annotations were linked to PGA domains and subdomains through the project trait-to-domain table. Taxonomic assignments were obtained from the accompanying species–family–primate-group table and were used to classify each species pair at family, superfamily, or order level (Kuderna et al., 2023).

Candidate columns were required to be numeric and to contain more than 20 non-missing observations. To limit discretized or quasi-categorical variables, we calculated value redundancy as 1 − (number of unique numeric values / number of observed values) and retained traits with redundancy below 0.10. The filter was applied independently to each column and generated both one standardized two-column input file per retained trait and a complete audit table. Under these criteria, 153 traits were retained for the main analysis.

#### 4.1.2 Phylogeny and name matching

Phylogenetic analyses used the Kuderna S4 primate tree distributed in the project as science.abn7829_data_s4.nex.tree (Kuderna et al., 2023). Tree manipulation was performed with ape (Paradis and Schliep, 2019). Species names in phenotypic tables and tree tip labels were standardized by removing apostrophes and replacing spaces with underscores. For each trait, finite observations were intersected with the tree, unmatched tips were removed, and the trait vector was reordered to the pruned tree. Traits with fewer than three matched species were not analysed. Consequently, every trait was evaluated on its own observed species set and pruned phylogeny; no missing phenotypes were imputed.

### 4.2 Analytical Phenotype Shift Score

The reference implementation of the score is provided in the phyloPSS R package (version [VERSION]; [DOI]). Pairwise scores are calculated with the exported function phylogenetic_shift_score(), which performs trait–tree matching, fits and selects between BM and OU models, computes S, Δ and T, and returns the complete ranked table of species pairs.

#### 4.2.1 Evolutionary model fitting and selection

For each trait, Brownian motion (BM) and single-optimum Ornstein–Uhlenbeck (OU) models were fitted with geiger::fitContinuous (Butler and King, 2004; Beaulieu et al., 2012; Pennell et al., 2014; https://cran.r-project.org/package=geiger) using one computational core. Model fits were required to return finite, positive diffusion variances; OU fits additionally required a finite positive attraction parameter α. The OU model was selected only when its AIC was more than two units lower than the BM AIC; otherwise BM was retained (Burnham and Anderson, 2002). This conservative rule treated BM as the default when model support was ambiguous.

#### 4.2.2 Model-based placement of pairwise differences

For every unordered species pair i,j, the observed absolute phenotypic difference was δᵢⱼ = |xᵢ − xⱼ|. Under either fitted Gaussian evolutionary model, the signed tip contrast has variance vᵢⱼ = Vᵢᵢ + Vⱼⱼ − 2Vᵢⱼ, where V is the model-specific phylogenetic covariance matrix. Because the fitted BM and homogeneous OU models assign a common expected value to tips, the absolute contrast follows a half-normal distribution. Its analytical placement score was therefore

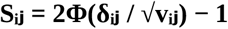

where Φ is the standard normal cumulative distribution function. Values near zero identify differences that are small relative to model expectation, whereas values near one identify differences near the upper end of the pair-specific expected distribution. Under BM, the contrast variance simplifies to σZdᵢⱼ, where dᵢⱼ is patristic distance. For OU, covariance was calculated from shared path length, patristic separation, σZ, and α on the possibly non-ultrametric pruned tree.

#### 4.2.3 Score construction

The absolute difference was normalized by the maximum pairwise difference observed for the trait, yielding Δᵢⱼ ∈ [0,1]. Patristic distance was normalized by the maximum pairwise distance on the trait-specific tree, yielding Tᵢⱼ ∈ (0,1]. The final score was

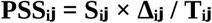

where S was taken from the selected evolutionary model. The complete table was sorted in decreasing PSS. PSS is a ranking index rather than a p-value, probability of convergence or divergence, or evolutionary-rate estimate. Division by T intentionally adds temporal weighting beyond the contribution of phylogenetic distance to the covariance underlying S.

### 4.3 Primate phenome workflow

#### 4.3.1 Nextflow implementation and empirical tails

The trait analysis was implemented in Nextflow DSL2 (Di Tommaso et al., 2017; https://www.nextflow.io/). The workflow sequentially filtered traits, calculated all pairwise PSS values, appended taxonomic classifications, selected empirical tails, generated cross-trait summaries, and ran trait-wise parametric bootstrap analyses. Pair classification used the family assignment when both species belonged to the same family, the superfamily/primate-group assignment when families differed but the higher group was shared, and order level otherwise. For a trait containing N pairs, the upper and lower tails each contained ceiling(0.01N) pairs selected from the decreasing PSS ranking. The workflow wrote raw, classified, tail, bootstrap, and summary outputs to separate reproducible directories.

#### 4.3.2 Trait-specific opportunity correction

Raw taxonomic tail composition was corrected for the pair types available for each trait. For taxonomic class k in trait t, opportunity share was pₜₖ = Nₜₖ/Nₜ, where Nₜ was the number of all tested pairs and Nₜₖ the number assigned to class k. Tail share qₜₖ was calculated from the selected upper or lower tail, and the primary plotted measure was qₜₖ − pₜₖ. Positive values indicate enrichment relative to opportunity and negative values indicate depletion. Classes with zero opportunity were recorded as not estimable rather than as zero enrichment. Exact hypergeometric reference intervals were calculated using the observed tail size; these were treated descriptively because species recur across pairs.

Phylogenetic depth was standardized within each trait by ranking every pair’s normalized patristic distance among all tested pairs. The opportunity-relative depth percentile was calculated as (average rank − 0.5)/Nₜ. For each tail, the median of these percentiles summarized whether selected pairs were concentrated among the shallowest or deepest comparisons actually available for that trait. Random-pair reference intervals were obtained from 2,000 samples without replacement per trait and tail size.

#### 4.3.3 Parametric calibration

Trait-wise parametric calibration used the same pruned tree and observed taxonomic pair universe as the empirical analysis. The model selected from the observed data was used as the generating process. Gaussian trait vectors were simulated from its fitted mean and covariance matrix. Each replicate was analysed twice: a conditional analysis retained the observed model parameters, whereas a full analysis refitted BM and OU, repeated the AIC-based model selection, recomputed PSS, and reselected the empirical tails. The Nextflow configuration specified 100 paired replicates per trait, a base seed of 20260812, and up to ten attempts for a valid replicate. Later depth-profile analyses in the project used larger replicate sets; the final manuscript should state the replicate number corresponding to the frozen analysis release.

For every replicate and tail we recorded opportunity-corrected family, superfamily, and order composition; median opportunity-relative depth; median S, Δ, T, and PSS; and model-selection stability. These simulations quantify the behaviour expected from the fitted evolutionary process, score geometry, trait coverage, and phylogenetic opportunity. They do not constitute a universal null model for all possible evolutionary mechanisms.

### 4.4 Relative brain-mass case study

#### 4.4.1 Phenotype definition and primate analysis

The genome–phenome case study used relative brain mass, defined as the signed residual from a mammal-wide quadratic regression of log brain mass on log body mass (Venditti et al., 2024). Residuals were calculated in the mammalian analysis and then transferred unchanged to primates, preserving the common mammalian allometric reference rather than refitting allometry within Primates. Exact phenotype–tree matching retained 129 primate species and 8,256 unordered pairs. BM and OU were refitted to this primate subset and PSS was calculated with the canonical model-selection and scoring procedure.

#### 4.4.2 Cercopithecidae-restricted analysis

A second PSS analysis was restricted to Cercopithecidae. The same relative-brain-mass residuals were copied without within-family renormalization, the Kuderna S4 tree was pruned to the matched family members, and BM and OU were refitted. The upper and lower 1% tails were calculated after ranking all within-family pairs. For the PSS-driven genome–phenome groups, a top-tail pair contributed to pool definition when at least one endpoint also appeared in a bottom-tail pair. Within each retained pair, the endpoint with higher relative brain mass was assigned to the foreground pool and the lower endpoint to the background pool; species were then deduplicated while preserving the pair-level evidence table.

### 4.5 Genome–phenome hypothesis benchmark

#### 4.5.1 Comparison strategies and pooled cycles

Nine predefined strategies were constructed to compare PSS-informed hypothesis formation with taxonomic and phenotype-only alternatives. Each strategy generated 100 unique cycles, each containing four foreground (FG) and four background (BG) species. Higher relative brain mass was consistently encoded as FG. Species retained fixed-side membership within an approach, which is required for pooled event reconstruction. A fixed seed (260821) and SHA-256-based ranking made cycle selection reproducible across software environments.

| No. | Strategy | Operational definition |
| --- | --- | --- |
| 1 | Family minima and maxima | Four families sampled per cycle; within each family the phenotypic maximum was FG and the minimum BG. |
| 2 | Primate-wide absolute tails | Four species independently sampled from high and low phenotype tails, with no repeated genus within either side. |
| 3 | Best PSS pair per genus | Four intact congeneric pairs sampled from the highest-PSS pair retained for each represented genus. |
| 4 | All congeneric top-1% PSS pairs | Four intact eligible pairs sampled from the global upper PSS tail after enforcing fixed-side membership. |
| 5 | Macaca–Papio–Trachypithecus PSS pairs | Top-1% PSS pairs restricted to the three prespecified genera. |
| 6 | Macaca–Papio PSS pairs | Top-1% PSS pairs restricted to the two focal Papionini genera. |
| 7 | Ranked endpoint-disjoint PSS matching | Thirteen non-overlapping directional PSS |
|  |  | pairs selected greedily; four intact pairs sampled per cycle. |
| 8 | Cercopithecidae PSS-driven pools | Nine FG and ten BG species defined by PSS; four species sampled independently from each fixed pool. |
| 9 | Cercopithecidae absolute tails | Upper and lower 10% within 55 family members defined six-species pools; four sampled per side. |

#### 4.5.2 Pooled CAAStools discovery

Each configuration file contained one cycle per row in the form cycle identifier, comma-separated FG species, and comma-separated BG species. A separate complete pool file defined fixed FG/BG membership for coherent event reconstruction and event denominators. For every alignment and strategy, the pooled-discovery implementation of CAAStools (Barteri et al., 2023; https://github.com/linudz/caastools) evaluated all 100 cycles. A residue was retained when at least one cycle was positive. Positive cycles at the same position were then combined into amino-acid-compatible events: a species was counted once within an event, whereas incompatible amino-acid signatures remained separate events.

Alignments were read in relaxed PHYLIP format. The positional hypergeometric prefilter was set to 0.05. Each 4-vs-4 hypothesis required at least three observed, non-gap species in both FG and BG; the maximum fraction of gaps per position was 0.5. The workflow emitted a legacy position table containing positive-cycle counts and identifiers and an event-level table used as the primary biological output. Cycles were treated as alternative realizations of the same hypothesis and not as statistically independent replicates.

#### 4.5.3 Cluster execution of pooled CAAStools

The pooled CAAStools benchmark was executed with a dedicated Nextflow DSL2 workflow (Di Tommaso et al., 2017) in the projectsu repository [ADD GITHUB REPOSITORY URL]. A tab-separated manifest linked every approach to its pooled configuration and complete pool. Nextflow combined this manifest with all amino-acid alignments, forming one independent job for each strategy–gene combination. Jobs called the bundled CAAStools pooled-discovery command and published position- and event-level outputs into approach-specific run directories.

Cluster execution used the SLURM workload manager (Yoo et al., 2003; https://slurm.schedmd.com/) through the Nextflow SLURM executor. The active configuration requested one CPU, 2 GB memory, and 30 min per gene-level pooled-discovery job, with a queue size of 100 and a submission-rate limit of one job every 10 s. The project Conda environment distributed in the supplementary/ directory of the phyloPSS repository (https://docs.conda.io/) was activated inside each task. The configured alignment glob targeted relaxed-PHYLIP files installed directly on the Correfoc cluster [Add font]. Failed or time-limited gene jobs were ignored so that remaining alignments could complete, whereas shared preparation failures terminated the workflow. Timestamped run identifiers separated outputs, and interrupted or extended graphs were continued using Nextflow’s resume mechanism and restricted one-row manifests when only a new strategy was to be scheduled.

### 4.6 CAAStools result consolidation and quality control

#### 4.6.1 Event and position filtering

The local meta-analysis consumed consolidated CAAStools outputs without rerunning discovery. Four finalized strategies were retained for the principal comparison: primate-wide phenotype extremes, family-level parallel pairs, Cercopithecidae PSS-driven groups, and Cercopithecidae-wide phenotype extremes. Primary events with nominal positional hypergeometric P < 0.05 were retained; no multiple-testing correction was applied at this discovery stage. A position was uniquely identified by gene and amino-acid coordinate. Duplicate primary records were resolved deterministically by lowest positional probability, then highest balanced support, then highest total support.

#### 4.6.2 Dense-position cluster filter

To reduce signals attributable to local alignment irregularities, indel boundaries, or extended sequence features, dense-position filtering was performed independently within each strategy and gene. A discovered position was discarded if it belonged to at least one interval containing three or more distinct positions, spanning at least three residues, and having density ≥0.70 hits per residue. Filtering was applied after selection of nominally significant primary positions and before any FG/BG support threshold. A gene was retained when at least one significant position survived and was classified as lost only when all its significant positions were removed.

#### 4.6.3 Species support and functional enrichment

Position-level FG and BG support were taken directly from the event table. Gene-level FG and BG support distributions used the largest support observed among significant positions, calculated separately for the two sides. Balanced support at a single position was defined as min(FG support, BG support); maxima observed at different residues were not combined. For functional analysis, genes were retained when at least one cluster-filtered significant position was supported by four or more FG and four or more BG species at that same coordinate.

Functional enrichment used g:Profiler (Kolberg et al., 2023; https://biit.cs.ut.ee/gprofiler/) with strategy-specific custom gene backgrounds derived exclusively from the corresponding consolidated CAAStools files. Human gene annotations were queried across Gene Ontology, KEGG, Reactome, WikiPathways, transcription-factor and microRNA targets, Human Protein Atlas, CORUM, and Human Phenotype Ontology sources (The Gene Ontology Consortium, 2023; Kanehisa et al., 2023; Gillespie et al., 2022; Martens et al., 2021; Uhlén et al., 2015; Tsitsiridis et al., 2023; Köhler et al., 2021; [Add font for regulatory-target sources]). Terms passing the g:SCS-adjusted threshold of 0.05 were considered significant. Fold enrichment was calculated as the fraction of query genes annotated to a term divided by the corresponding fraction in the effective custom background. Strategy comparison emphasized the joint relationship between query-set selectivity, enrichment strength, and the breadth of significant functional terms rather than raw discovery counts alone.

### 4.7 RERconverge branch

The supplementary/ directory of the phyloPSS repository also contains an independent RERconverge branch (https://github.com/nclark-lab/RERconverge; Kowalczyk et al., 2019) for testing associations between relative evolutionary rates and the same binary phenotype configurations. Gene trees are constructed on the reference topology with gene-specific branch lengths, assembled into a multi-gene manifest, and analysed jointly because relative-rate estimation requires multiple genes. The branch is scheduled independently of CAAStools. At the time represented by the current supplementary validation materials, however, the RERconverge workflow was planned but no consolidated result set was available. Its detailed inferential settings and final comparison metrics should therefore be added only after the analysis is frozen.

### 4.8 Reproducibility and provenance

All empirical, bootstrap, case-study, pooled-discovery, and meta-analysis stages were implemented as scripted R, Python, and Nextflow workflows distributed under supplementary/ in the phyloPSS repository. Inputs, generated configuration manifests, random seeds, checksums, model-fit tables, and audit tables were retained alongside the analyses. The original uncorrected taxonomic summary and superseded exploratory outputs were preserved in dedicated provenance locations rather than overwritten. The manuscript release should archive the frozen code, configuration manifests, software environments, and final result tables and should report the exact phyloPSS version and commit used for score calculation, together with the exact versions of R, Python, Nextflow, ape, geiger, CAAStools, g:Profiler data, and RERconverge.

## Author contributions

F.B.: Conceptualization, Methodology, Software, Validation, Formal analysis, Investigation, Data curation, Visualization, Writing – original draft, Writing – review & editing, and Project administration. A.N.: Conceptualization, Methodology, Supervision, and Writing – review & editing. O.E.C.: Conceptualization, Methodology, Supervision, and Writing – review & editing. All authors contributed to the interpretation of the results and approved the final manuscript.

