## Supplementary Notes and Figures for "Phenotype shift scores reveal the scale and phylogenetic structure of phenotypic differentiation in primates"

#### Supplementary Note 1. Positioning Phenotype Shift Scores among phylogenetic approaches to phenotypic differentiation

This focused comparison defines the analytical space occupied by Phenotype Shift Scores (PSS). It is not intended as a systematic review. Instead, it contrasts the principal questions, units of analysis and inferential outputs of methods that quantify phylogenetically structured phenotypic change, similarity, convergence or evolutionary regime shifts.

##### 1.1. Distinct analytical meanings of phenotypic change

Methods described as detecting phenotypic shifts often address different biological objects. A shift may refer to a standardized difference between sister lineages, an unusual contrast between extant species, a reconstructed increase in similarity, a change on a particular branch, or a transition between evolutionary regimes. These objects are not interchangeable. In particular, an exceptional difference between two terminal species does not by itself identify the branch on which change occurred, demonstrate a change in evolutionary rate, or establish adaptation.

The relevant methods also differ in search scope. Some test focal taxa or regimes specified in advance, whereas others explore many nodes, branches or clades. Their units may be terminal pairs, internal-node contrasts, focal groups, reconstructed lineages, branches or adaptive regimes. They further differ in their dependence on ancestral-state reconstruction, treatment of univariate and multivariate traits, and ability to evaluate similarity and differentiation symmetrically. These distinctions provide the framework for positioning PSS.

##### 1.2. Existing approaches to contrasts, convergence and evolutionary shifts

Phylogenetic independent contrasts (PIC) transform species values into n - 1 contrasts between sister descendant lineages, standardized by the variance expected from their branch lengths under Brownian motion (Felsenstein, 1985; Garland et al., 1992). PIC and PSS therefore share the principle that a phenotypic difference should be interpreted relative to model-predicted evolutionary variance. Their purposes and units differ: PIC generates nodal contrasts for association analyses, whereas PSS preserves the identities of arbitrary terminal pairs and evaluates the complete pairwise matrix.

Distance-based methods ask whether phenotypic resemblance is surprising given phylogenetic separation, quantify multivariate occupation of morphospace, or measure the compactness and distinctiveness of a predefined focal group (Stayton, 2006; Arbuckle et al., 2014). Probabilistic distances under continuous-trait models provide a related model-aware perspective but compare phylogenetic models and their predictions rather than screening terminal species pairs (Adams et al., 2021). These approaches are important conceptual antecedents, although terminal resemblance alone cannot distinguish acquired convergence from prolonged conservatism.

Historical convergence measures address that distinction more directly. Stayton's C1-C4 statistics quantify the proportion or magnitude of historical phenotypic distance closed by focal lineages, while C5 counts independent entries into a region of morphospace (Stayton, 2015). The time-synchronized Ct1-Ct4 measures reduce biases caused by comparing reconstructed lineage positions from different times and can identify increasing similarity or differentiation (Grossnickle et al., 2024). The Wheatsheaf index and search.conv instead evaluate the strength of similarity within focal groups or among clades (Arbuckle et al., 2014; Castiglione et al., 2019). These methods provide historical, geometric or clade-level information that terminal-pair scores do not contain.

A separate family of methods estimates changes in evolutionary process. SURFACE searches for branches that shift towards Ornstein-Uhlenbeck regimes and subsequently identifies regimes that can share an optimum (Ingram and Mahler, 2013). l1ou, bayou and phyloEM use penalized, Bayesian or expectation-maximization approaches to estimate the location and magnitude of adaptive shifts (Uyeda and Harmon, 2014; Khabbazian et al., 2016; Bastide et al., 2018). Their outputs concern branches, regimes and adaptive landscapes rather than the relative extremeness of every terminal contrast.

#### 1.3. The unresolved problem of exhaustive terminal-pair discovery

None of these distinctions implies a hierarchy among methods: each is suited to a different question. They nevertheless leave a practical gap when the aim is exploratory. A researcher may wish to identify which extant species pairs are more similar or more different than expected, compare pairs separated by very different evolutionary distances, preserve the identities of the species involved, and then locate the resulting contrasts across a phylogeny. Performing this operation exhaustively also creates a specific statistical structure: the number of pairs grows rapidly with taxon sampling, each species contributes to many observations, and the opportunities available at different phylogenetic depths are unequal.

Consequently, an all-pairs framework must separate ranking from inference. Percentile tails can define operational sets of exceptional contrasts, but their members are neither independent observations nor automatically significant evolutionary events. Claims about aggregate depth or nodal concentration require null models that reproduce the fitted evolutionary process, the reuse of species across pairs, model refitting and the available pairwise opportunity.

#### 1.4. The methodological position and limits of PSS

PSS occupies this discovery layer. For each terminal pair, it converts the observed absolute phenotypic difference into its percentile under a continuous-trait evolutionary model fitted to the trait-specific pruned phylogeny. The common percentile scale permits contrasts at different patristic distances to be ranked together. The lower tail identifies pairs that are unusually similar under the model, whereas the upper tail identifies pairs that are unusually different. Because species identities are retained, selected pairs can be summarized by evolutionary depth, taxonomic composition or most recent common ancestor and can be used to construct targeted genome-phenome comparisons.

PSS is therefore complementary to, rather than a replacement for, established phylogenetic comparative methods. It does not reconstruct ancestral trajectories, locate a causal change on a branch, estimate adaptive optima or selection strength, or demonstrate historical convergence. PIC remains appropriate for independent contrasts in trait-association analyses; Ct measures are more informative when the question is whether focal lineages became more similar through time; and regime-shift methods are required when the aim is to infer changes in evolutionary process. PSS instead provides a model-calibrated and bidirectional map from which those focused analyses can begin.

In the present framework, empirical percentile tails are treated as discovery sets. Their phylogenetic-depth distributions are evaluated with a full parametric bootstrap in which traits are simulated under the fitted Brownian-motion or Ornstein-Uhlenbeck model, models are refitted and reselected, and the complete PSS workflow is repeated. Pairwise opportunity is incorporated when interpreting taxonomic depth and node-level concentration. This combination distinguishes the descriptive localization of terminal contrasts from statistical evidence that their aggregate phylogenetic structure departs from the fitted null expectation.

Methodological conclusion. Existing methods standardize nodal contrasts, reconstruct convergence and locate changes among evolutionary regimes. PSS complements them by providing an exhaustive, model-calibrated and bidirectional map of exceptional phenotypic differentiation among terminal species pairs.

**Supplementary Table 1. Comparison of methods relevant to exceptional phenotypic differentiation**

*The table compares primary analytical targets rather than ranking methods. 'Global' indicates that the search domain spans a substantial part of the phylogeny; it does not imply that the methods use the same unit of analysis or inferential procedure.*

| Method | Unit | Primary question | Search | Ancestors | Direction | Calibration / multiplicity | Relation to PSS |
|---|---|---|---|---|---|---|---|
| PIC | Internal-node contrast | How can trait associations be analysed without treating species as independent? | Full-tree transform | Recursive nodal values | Signed contrast | Independent under adequate BM and branches; no all-pairs family | Shares variance standardization, but not arbitrary terminal-pair discovery |
| Phenotypic / phylogenetic distance ratios | Focal pair or group | Is resemblance surprising given evolutionary separation? | Usually focal | Not required | Mostly similarity | Simulation or randomization varies by implementation | Conceptual antecedent; PSS supplies pair-specific model percentiles and both tails |
| Probabilistic continuous-trait distances | Phylogenetic model or tree | How different are phylogeny-aware models and their predictions? | Model-level | Not required | Not applicable | Probability-distribution distances; not a terminal-pair testing family | Model-aware but addresses distances among models, not among species pairs |
| Wheatsheaf | A priori focal group | How compact and distinctive is a proposed convergent group? | Confirmatory | Not required | Similarity | Bootstrap / permutation | Quantifies a known group; PSS can discover candidate pairs or members |
| C1-C4 | Focal lineages | How much historical phenotypic distance was closed? | Confirmatory | Required | Convergence | BM simulations for focal hypotheses | Adds reconstructed history; PSS avoids ancestors and also evaluates differentiation |
| C5 | Morphospace region | How many independent lineages entered a phenotype region? | Focused | Required | Independent origins | Simulation; depends on region definition | Counts origins rather than ranking terminal extremeness |
| Ct1-Ct4 | Focal lineage pair / clades | Did lineages become more similar or different at comparable times? | Confirmatory | Required | Both | BM simulations for focal hypotheses | Natural historical follow-up to PSS; same-data follow-up is conditional |
| search.conv | Node / clade pair | Are clades or states more similar than expected? | Focused or automatic | Partial | Similarity / direction | Randomization and node-distance filtering | Adds clade-level direction; not an exhaustive terminal-pair scan |
| SURFACE | Branch and OU regime | Did lineages shift towards shared adaptive optima? | Global | Regimes inferred | Shared optima | Stepwise AICc model selection | Infers process and regimes; PSS maps terminal contrasts |
| l1ou / bayou / phyloEM | Branch and shift configuration | Where and how large are adaptive shifts? | Global | Latent regimes | Shift / regime | Penalized, Bayesian or EM inference | Process models for explaining structures highlighted by PSS |
| PSS | Terminal species pair | Which terminal differences are exceptional under a fitted evolutionary model? | All pairs | Not required | Similarity and differentiation | Percentile ranking; opportunity correction and full parametric bootstrap for aggregate patterns | Discovery and localization layer; does not infer historical process |

#### Supplementary Note 2. Analytical construction and empirical behaviour of the Phenotype Shift Score.

This note provides the analytical derivation and empirical interpretation of the Phenotype Shift Score (PSS), using allometry-adjusted brain mass in primates as a worked example. The phenotype is the signed residual from the mammal-wide brain-body relationship and is later used to localize relative-brain-mass differentiation and construct genome-phenome comparisons. Here, the comparison between *Homo sapiens* and *Gorilla gorilla* illustrates how an observed trait difference is converted into a model-calibrated, phylogeny-wide rank. The example is explanatory rather than inferential: its Upper 1% membership is a prioritization outcome, not a declaration of statistical significance.

##### 2.1. Inferential target and notation

For each continuous trait, observed species are matched to a phylogeny, the tree is pruned to those species and every unordered terminal pair is evaluated. For species i and j, xi and xj denote observed trait values, deltaij = |xi - xj| is their absolute phenotypic difference and dij is their patristic distance. PSS combines three related quantities: the placement of deltaij under a fitted evolutionary model, its magnitude relative to all observed pairwise differences and the phylogenetic separation across which it occurs.

The three quantities are intentionally not independent. The observed difference enters both model placement and relative magnitude, while phylogenetic separation contributes to the model-predicted variance and appears again as an explicit denominator. PSS should therefore be interpreted as a designed ranking index that reinforces concordant signals, not as a combination of independent tests.

##### 2.2. Analytical derivation of the model-based placement score

*2.2.1 Gaussian tip distribution and pairwise contrasts*

Let model M predict a multivariate Gaussian distribution for the vector of trait values at the tips:

$$X \sim \mathrm{N}(mu^{(\mathrm{M})}, V^{(\mathrm{M})}). \qquad [\mathrm{S1}]$$

A contrast vector aij containing +1 for species i, -1 for species j and zero elsewhere extracts the signed terminal contrast:

$$D_{ij} = X_i - X_j = a_{ij}^T X. \qquad [\mathrm{S2}]$$

Because a linear contrast of a multivariate normal vector is normal, Dij has mean mij(M) and variance vij(M). The covariance term is essential because the two species share evolutionary history:

$$D_{ij} \sim \mathrm{N}(\mathrm{m}_{ij}^{(\mathrm{M})}, v_{ij}^{(\mathrm{M})}), \quad v_{ij}^{(\mathrm{M})} = V_{ii}^{(\mathrm{M})} + V_{jj}^{(\mathrm{M})} - 2V_{ij}^{(\mathrm{M})}. \quad [\mathrm{S3}]$$

*2.2.2 From the signed contrast to S*

The homogeneous BM and single-optimum OU models used here assign a common expected value to the two tips, so mij(M) = 0. The signed contrast is therefore centred on zero and its absolute value follows a half-normal distribution with scale equal to the square root of vij(M). Evaluating the half-normal cumulative distribution at the observed difference gives

$$S_{ij}^{(\mathrm{M})} = P_M(|\mathrm{D}_{ij}| <= \mathrm{delta}_{ij}) = 2\ \mathrm{Phi}(\mathrm{delta}_{ij} / \mathrm{sqrt}(\mathrm{v}_{ij}^{(\mathrm{M})})) - 1. \qquad [\mathrm{S4}]$$

S is the percentile occupied by the observation in the pair-specific theoretical distribution of absolute differences. An S value of 0.95 means that 95% of absolute differences predicted for that pair are no greater than the observed one. It does not mean that divergence occurred with probability 0.95, that the selected model has probability 0.95 or that the pair passes an error-controlled statistical test. If a future evolutionary model assigns different expected values or optima to the two tips, the absolute contrast becomes folded-normal and equation [S4] must be replaced by the corresponding folded-normal cumulative distribution.

*2.2.3 BM and OU pairwise variance*

Under homogeneous Brownian motion, the expected variance of a terminal contrast increases linearly with patristic distance (Felsenstein, 1985):

$$v_{ij}^{(\mathrm{BM})} = \mathrm{sigma}_{BM}^2\ d_{ij}. \qquad [\mathrm{S5}]$$

For a homogeneous single-optimum OU process on an ultrametric tree, attraction towards a common optimum causes pairwise variance to approach a finite limit rather than increase without bound (Hansen, 1997; Butler and King, 2004):

$$v_{ij}^{(OU)} = (\text{sigma}_{OU}^2 / \text{alpha})[1 - \exp(-\text{alpha } d_{ij})]. \qquad [S6]$$

Both models are fitted once to the complete matched trait vector and pruned tree. Their fitted covariance matrices determine model-specific placement values for every pair, and the selected model supplies the S value used in the final score. A consistent branch-length scale is required for the fitted rate, OU attraction parameter and pairwise distances.

*2.2.4 Relative magnitude, relative distance and the composite score*

S locates a difference relative to the evolutionary expectation for that pair but does not retain its magnitude relative to the range observed in the analysed dataset. The absolute difference and patristic distance are therefore normalized by their maxima across all available pairs:

$$\text{Delta } x_{rel,ij} = \text{delta}_{ij} / \max_{k<l}(\text{delta}_{kl}), \quad d_{rel,ij} = d_{ij} / \max_{k<l}(d_{kl}). \quad [S7]$$

The final score multiplies model placement by relative magnitude and divides by relative distance:

$$PSS_{ij} = S_{ij}(\text{Delta } x_{rel,ij} / d_{rel,ij}). \qquad [S8]$$

Phylogenetic distance consequently enters twice for two distinct purposes. It first determines how much evolutionary variance is expected under the fitted model and then provides an explicit additional weighting that favours disparity concentrated over short separation. This is an intentional design choice. PSS is dimensionless but not bounded above by one, and its values are most directly comparable within analyses sharing the same taxa, tree, trait definition and preprocessing.

Equations [S1–S8] are implemented in the phyloPSS R package through phylogenetic_shift_score(), which returns the fitted models, model-comparison summary and complete ranked table of species pairs.

#### 2.3. Worked calculation for Homo sapiens and Gorilla gorilla

The focal analysis retained the mammal-wide allometry-adjusted brain-mass residual for 129 primates matched to the Kuderna et al. (2023) phylogeny, producing 8,256 unordered pairs. Homo sapiens and Gorilla gorilla had residual values of 2.9245501989 and 1.6262969317. Their observed contrast and distance were

$$\text{delta}_{HG} = |2.9245501989 - 1.6262969317| = 1.2982532672, \quad d_{HG} = 20.238906202. \qquad [S9]$$

The maximum phenotypic difference and maximum patristic distance across the same 8,256-pair universe were 2.9970907694 and 121.413057327. Consequently,

$$\text{Delta } x_{rel,HG} = 1.2982532672 / 2.9970907694 = 0.4331711540, \quad d_{rel,HG} = 20.238906202 / 121.413057327 = 0.1666946426. \quad [S10]$$

The archived model-fit table reports information-criterion values of 21.8115 for BM and 19.5615 for OU, giving OU a 2.2500-unit advantage. The selected OU fit had sigma2 = 0.0093325870 and alpha = 0.0153271152. At the focal distance, equation [S6] gives an expected pairwise variance of 0.1623936387 and a standard deviation of 0.4029809408. The standardized absolute contrast and placement were

$$z_{HG} = 1.2982532672 / 0.4029809408 = 3.221624488, \quad S_{HG} = 2 \text{ Phi}(3.221624488) - 1 = 0.9987253393. \quad [S11]$$

Using the unrounded components, the complete score was

$$PSS_{HG} = 0.9987253393(0.4331711540 / 0.1666946426) = 2.5952784138. \qquad [S12]$$

This value ranked 22nd among 8,256 comparisons. Because the Upper 1% contained the 83 highest-ranking pairs, the focal comparison entered that prioritization set. Its placement illustrates that a moderately scaled observed disparity can rank highly when it is both extreme under the selected evolutionary model and concentrated over relatively limited phylogenetic separation.

#### 2.4. Empirical component behaviour in Supplementary Figure 1

##### *2.4.1 Analytical placement and model-specific variance*

Panel a displays the exact monotonic transformation from standardized absolute contrast to S. The curve rises rapidly and asymptotically approaches one; the focal z value of 3.2216 maps to S = 0.9987. The near-saturation of S is therefore an analytical consequence of the contrast being more than three expected standard deviations from zero, rather than an empirical threshold imposed after observing the ranking.

Panel b contrasts the distance dependence of the two fitted models. BM variance increases linearly, whereas OU variance bends towards its finite asymptote. At the focal distance of 20.2389, the fitted BM and OU contrast variances are similar (0.16434 and 0.16239), but their trajectories separate progressively at greater distances. This geometry explains why a model-specific covariance is required: the same raw disparity can occupy a different theoretical percentile under BM and OU, especially for widely separated species.

##### *2.4.2 Empirical component geometry*

Panel c places the analytical relationship in empirical context using male body mass, selected as BM and represented by 9,730 pairs from 140 species, and mean precipitation, selected as OU and represented by 7,626 pairs from 124 species. S increases with relative phenotypic difference in both examples because both quantities contain the observed disparity. Their relation is not one-to-one, however, because the pair-specific variance inside S also depends on phylogenetic covariance. The Upper 1% sets occupy combinations of large S and large disparity-to-distance ratio, whereas Lower 1% pairs cluster near zero for both terms. The component plots are descriptive geometry, not r egressions among independent variables: every species contributes to many pairwise observations and the displayed components are mathematically coupled.

##### *2.4.3 Algorithmically matched pair controls*

Panel d compares the focal result with two controls selected by declared numerical rules before species identities were inspected. Macaca maura and Avahi laniger had nearly the same relative phenotypic difference as the focal pair (0.43318 versus 0.43317) but the maximum relative distance (1.000). Their S remained high at 0.92979, yet the long separation reduced PSS to 0.40277 and rank 1,064. Pan paniscus and Gorilla gorilla had the same relative distance as the focal pair (0.16669) but a much smaller relative difference (0.06686). Their S was 0.38097, PSS was 0.15279 and the pair ranked 4,036th. These controls show directly that the focal score is not recoverable from raw disparity or distance alone.

##### *2.4.4 Component ablation*

Panel e recalculates rankings after removing one component at a time. Omitting S retained a Spearman rank correlation of 0.9676 with complete PSS and moved the focal pair from rank 22 to 26, but the Jaccard overlap between Upper 1% sets fell to 0.581. Omitting relative phenotypic magnitude reduced rank correlation to 0.8061, Upper 1% overlap to 0.456 and the focal rank to 124. Omitting the explicit distance denominator retained a rank correlation of 0.9045 but only 0.099 Upper 1% overlap and moved the focal pair to rank 372. The high global correlations therefore do not imply that the same pairs occupy the prioritization tail. The explicit distance weighting is particularly influential for Upper 1% membership, while S and phenotypic magnitude also materially change which pairs are retained.

##### *2.4.5 Sensitivity to alternative tail widths*

Panel f shows the empirical score boundaries produced by four prespecified tail widths. The 0.5%, 1%, 2.5% and 5% definitions selected 42, 83, 207 and 413 pairs per tail. Their Upper boundaries were 1.52154, 1.13907, 0.81549 and 0.56589, whereas their Lower boundaries were 1.831 x 10^-5, 7.019 x 10^-5, 3.756 x 10^-4 and 1.529 x 10^-3. The focal PSS of 2.5953 remained above every Upper boundary. Changing tail width alters the number of prioritized comparisons and the empirical cutoff but does not convert the tail into a null-model significance test.

##### *2.5. Interpretation and limitations*

Together, the six panels establish that PSS is not a relabelled raw difference, a simple inverse-distance score or a probability of evolutionary change. S calibrates disparity against a model- and pair-specific contrast distribution; relative phenotypic magnitude retains the position of that disparity within the observed trait range; and relative distance adds an explicit weighting for phylogenetic concentration. The matched controls and component ablations show that each ingredient can alter pair ranks and tail membership.

The same construction also imposes interpretive limits. Both maxima used for normalization depend on taxon sampling. S and relative magnitude share the same observed difference, while distance contributes to both S and the final denominator. Very short branches can be amplified, pairs are statistically dependent because species recur across comparisons and an extreme terminal contrast does not identify the historical branch on which change occurred. Consequently, raw PSS values should not be compared across unrelated datasets without examining their components, and percentile tails should be used for prioritization and downstream hypothesis construction rather than as direct evidence of convergence, divergence or altered evolutionary rate.

#### Supplementary Table 1. Exact computational audit of the worked example

*a, Focal Homo sapiens-Gorilla gorilla calculation. Values retain the precision stored in the frozen Figure S1 audit. The information criterion is labelled AIC in the source table.*

| Stage | Quantity | Symbol | Exact value | Interpretation |
|---|---|---|---|---|
| Input | H. sapiens trait value | xH | 2.92455019888865 | Mammal-wide allometry-adjusted brain mass |
| Input | G. gorilla trait value | xG | 1.62629693168425 | Mammal-wide allometry-adjusted brain mass |
| Observed | Absolute difference | deltaHG | 1.29825326720440 | Absolute terminal disparity |
| Observed | Patristic distance | dHG | 20.238906202 | Distance on the pruned primate tree |
| Scaling | Maximum difference | deltamax | 2.99709076939713 | Maximum across all 8,256 pairs |
| Scaling | Maximum distance | dmax | 121.413057327 | Maximum across all 8,256 pairs |
| Component | Relative difference | Delta xrel | 0.433171153993960 | deltaHG / deltamax |
| Component | Relative distance | drel | 0.166694642632142 | dHG / dmax |
| Model fit | BM information criterion | AICBM | 21.8115112354088 | Source-table label: AIC |
| Model fit | OU information criterion | AICOU | 19.5615356992570 | OU advantage = 2.24997553615178 |
| Model fit | Selected model | M* | OU | Used to calculate final S |
| Model fit | OU diffusion variance | sigma2OU | 0.00933258695100565 | Fitted on 129 primates |
| Model fit | OU attraction strength | alpha | 0.0153271152010042 | Fitted on the same branch-length scale |
| Calibration | Expected pairwise variance | vHG(OU) | 0.162393638680482 | Equation [S6] at dHG |
| Calibration | Standardized contrast | zHG | 3.22162448799061 | deltaHG / sqrt(vHG) |
| Calibration | Model placement | SHG | 0.998725339252273 | Half-normal cumulative placement |
| Output | Phenotype Shift Score | PSSHG | 2.59527841384567 | Complete score using unrounded values |
| Output | Rank | rank | 22 of 8,256 | Within the 83-pair |

| Stage | Quantity | Symbol | Exact value | Interpretation |
|---|---|---|---|---|
| | | | | Upper 1% |

#### Supplementary Table 1b. Algorithmically matched pair controls

*Controls were selected by fixed numerical rules before species identities were inspected. Delta rel, relative phenotypic difference; d rel, relative patristic distance.*

| Pair | Selection rule | Delta rel | d rel | S | PSS | Rank | Set |
|---|---|---|---|---|---|---|---|
| H. sapiens-G. gorilla | Focal pair | 0.433171 | 0.166695 | 0.998725 | 2.595278 | 22 | Upper 1% |
| M. maura-A. laniger | Closest phenotype among pairs with d rel >= 2 x focal | 0.433182 | 1.000000 | 0.929787 | 0.402767 | 1,064 | Other |
| P. paniscus-G. gorilla | Closest distance among pairs with Delta rel <= 0.5 x focal | 0.066855 | 0.166695 | 0.380967 | 0.152792 | 4,036 | Other |

#### 2.2 Extreme phenotypic differentiation occupies trait-specific phylogenetic depths in primates

Because the Phenotype Shift Score (PSS) favours phenotypic differences concentrated across short relative phylogenetic distances, the location of its empirical tails cannot by itself distinguish biological structure from score behaviour. We therefore evaluated PSS across 153 continuous traits retained from the Primate Genotype-Phenotype Archive (PGA; Valenzuela et al., 2026). These traits were selected from 236 numeric candidates with at least three observations by requiring more than 20 non-missing observations and less than 10% redundancy. The retained set comprised 3 behavioural, 5 ecological, 14 life-history, 89 morphological and 42 physiological traits. After matching trait records to the S4 primate phylogeny, coverage ranged from 19 to 140 species and from 171 to 9,730 unordered pairs per trait (median, 36 species and 630 pairs). Family coverage remained strongly trait specific, making separate tree pruning and model fitting essential for every analysis (Supplementary Fig. 2a-d and Supplementary Table 2). For each retained trait, Brownian-motion (BM) and Ornstein-Uhlenbeck (OU) models were compared, all available species pairs were scored, and the Upper and Lower 1% of the trait-specific PSS distribution were defined as candidate sets rather than intrinsically significant classes.

We first expressed each tail's phylogenetic depth relative to the pair opportunities on its pruned tree (Fig. 2a). A percentile of 0% identifies the shallowest available comparisons, 50% the median depth and 100% the deepest comparisons. Across traits, the median of the trait-specific Upper 1% depth medians was 2.96% (interquartile range, 1.59-13.33%), compared with 55.93% (39.91-63.33%) for the Lower 1%. High-ranking PSS pairs were therefore generally concentrated among relatively shallow comparisons. This tendency is expected from the inverse distance component of PSS and should not be interpreted as a universal biological preference for differentiation among close relatives; the raw depth distribution combines trait structure with the geometry of the score.

The comparisons available at each taxonomic depth also depended on trait coverage. Across the 153 retained traits, order-wide comparisons between major primate groups were estimable for 117 traits, whereas between-family and within-family, between-genus comparisons were available for all traits and within-genus comparisons for 147. The 36 traits lacking order-wide opportunity were concentrated in body-mass, longevity and reproduction measurements, while the six traits lacking within-genus opportunity were brain measurements (Supplementary Fig. 3a,b,d and Supplementary Table 3). We therefore measured taxonomic excess as the fraction of tail pairs at a focal level minus the corresponding fraction among all tested pairs for the same trait; zero denotes proportional representation (Fig. 2b; Supplementary Fig. 3c). Tail excess varied substantially within domains and subdomains, and no primary PGA domain was uniformly associated with one phylogenetic scale. For example, the Body mass Upper 1% was enriched within families by 30.6 percentage points and depleted order-wide by 36.2 points, whereas mean annual precipitation showed corresponding excesses of +82.0 and -70.9 points. Opportunity adjustment thus made traits with different sampling geometries comparable, but it remained descriptive and did not by itself demonstrate departure from the fitted evolutionary process.

We tested for such departures with 1,000 full parametric-bootstrap replicates per trait. In every replicate, phenotypes were simulated under the selected BM or OU model, both models were refitted and reselected, all pairwise PSS values were recalculated, and the same Upper and Lower 1% definitions were reapplied. The observed cumulative depth profiles were compared with the bootstrap median and simultaneous 95% envelope, and directional signed-area probabilities were corrected across traits using the Benjamini-Hochberg procedure. BM was selected for 92 traits and OU for 61; model-selection stability across the full refits ranged from 0.497 to 1.000, with 118 traits retaining the generating model in at least 90% of replicates (Supplementary Fig. 4a-c and Supplementary Table 4). At a false-discovery rate of 0.10, 20 Upper 1% profiles showed supported deepward redistribution and two Lower 1% profiles showed supported shallowward redistribution (Supplementary Fig. 4d,e). The deepward set comprised 11 body-mass measurements, five brain measurements, three biometabolite traits and territoriality. Related body-mass and brain definitions represent robustness across correlated measurements rather than independent biological replications. The inferential result is the deviation of a complete depth profile from its refitted null, not membership of a pair in an empirical 1% tail.

Body mass and mean annual precipitation illustrate why raw taxonomic or nodal localization required model calibration (Fig. 2c,d). Body mass (Smaers et al., 2021) included 136 species, 9,180 pairs and 92 Upper 1% pairs, with BM selected and retained in 95.4% of bootstrap refits. Its observed cumulative profile fell below the refitted-null profile over restricted depth intervals, yielding a signed area of -0.126 and a directional FDR-adjusted $q = 0.056$. By contrast, mean annual precipitation (Kamilar and Cooper, 2013) included 124 species, 7,626 pairs and 77 Upper 1% pairs. OU was selected in every bootstrap replicate, its signed area was -0.0019 ($q = 0.900$), and the observed profile never crossed the simultaneous envelope. Body mass therefore exhibited a deepward departure from the depth distribution generated by its fitted BM process, whereas the more strongly shallow precipitation profile was compatible with its fitted OU expectation.

The circular MRCA maps localized these pairwise configurations without converting them into branch- or node-level tests. Each Upper 1% pair was assigned once to its most recent common ancestor, and node size and colour summarized pair counts and opportunity-normalized concentration. Deeper internal separations accumulated more high-ranking body-mass pairs than expected from the complete fitted null profile, whereas precipitation pairs remained concentrated mainly at shallow nodes within the range expected under OU. Thus, the relevant contrast was not simply that one trait produced deep pairs and another shallow pairs. The phenome-wide analysis identified trait-specific redistributions of extreme differentiation beyond both taxonomic opportunity and the intrinsic geometry of PSS, while retaining the MRCA maps as descriptive localizations of the extant species pairs that contributed to those departures.

#### Supplementary Note 3. Sampling geometry, taxonomic opportunity and null calibration of the primate phenome-wide PSS analysis.

This note documents the construction and inferential calibration of the phenome-wide primate analysis supporting Main Results Section 2.2. The analysis was designed around three distinct layers. First, trait filtering and phylogenetic matching determine which species and species pairs can contribute to each trait. Second, opportunity-adjusted summaries describe where Upper and Lower 1% PSS pairs occur relative to the comparisons actually available. Third, a full parametric bootstrap asks whether an observed phylogenetic-depth profile departs from the profile generated when the complete PSS workflow is repeated under the fitted evolutionary model. Supplementary Figures 2-4 correspond to these three layers. Keeping them separate prevents incomplete trait coverage, the built-in distance weighting of PSS and null-model departure from being interpreted as the same phenomenon.

##### 3.1. Scope and analytical hierarchy

For every retained continuous trait, observations were matched to the S4 primate phylogeny and the tree was pruned to the matched species. BM and single-optimum OU models were fitted to that trait-specific dataset, and every unordered species pair on the pruned tree was assigned a PSS. The Upper and Lower 1% of the resulting empirical distribution were used as operational prioritization sets. These tails identify the highest- and lowest-ranking comparisons within a trait, but they do not constitute pairwise hypothesis tests and their members are not statistically independent because each species occurs in many pairs.

The three supplementary figures answer progressively narrower questions. Supplementary Figure 2 asks which traits and taxa enter the analysis. Supplementary Figure 3 asks which phylogenetic and taxonomic comparisons are available and whether a tail contains more or fewer pairs at a given level than expected from that opportunity. Supplementary Figure 4 asks whether the complete observed depth profile is displaced relative to a null distribution that includes simulation, model refitting, model reselection, score recalculation and tail reselection. Only the last layer supports inferential claims about a trait-level redistribution of extreme phenotypic differentiation.

##### 3.2. Trait retention and matched phylogenetic coverage

The starting audit contained 236 numeric PGA traits with at least three observations. Two prespecified filters were applied independently: more than 20 non-missing observations and less than 10% redundancy, where redundancy was defined as one minus the proportion of unique numeric values. Of the 236 candidates, 222 passed the observation criterion, 165 passed the redundancy criterion and 153 passed both (Supplementary Fig. 2a). The retained dataset comprised 3 Behaviour, 5 Ecology, 14 Life history, 89 Morphology and 42 Physiology traits distributed across the PGA subdomains shown in Supplementary Figure 2b. The filters therefore removed both sparsely sampled traits and variables whose nominally continuous values were dominated by repeated categories or coarse coding.

Observation counts in the source table were not treated as equivalent to phylogenetic coverage. Species names were matched to the tree after filtering, and only matched species determined the universe of tested pairs. Matched coverage ranged from 19 to 140 species and from 171 to 9,730 unordered pairs per trait; the median was 36 species and 630 pairs (Supplementary Fig. 2c). Because pair number grows quadratically with species number, apparently moderate differences in matched coverage produced large differences in the amount of pairwise opportunity and in the numerical resolution of a 1% tail.

Coverage was also taxonomically uneven. The family-by-trait matrix in Supplementary Figure 2d summarizes the proportion of classified S4 species represented in 16 families belonging to five major primate groups. Some traits spanned most major lineages, whereas others were densely sampled within only one or a few families. Consequently, the absence of deep or shallow tail pairs could reflect a missing opportunity rather than a biological exclusion. This is why each trait was analysed on its own pruned tree and why comparisons of raw pair counts across traits were avoided.

#### 3.3. Taxonomic opportunity and estimability

Taxonomic opportunity was partitioned into four mutually exclusive categories: comparisons between major primate groups, comparisons between families within the same major group, comparisons between genera within a family and comparisons within a genus. The broader term order-wide refers to the first category, whereas within-family summaries combine the two shallowest categories when used in Main Figure 2. The categories describe the taxonomy of extant species pairs; they do not identify the historical branch on which phenotypic change occurred.

Order-wide opportunity was estimable for 117 of 153 traits. The 36 non-estimable traits comprised 23 body-mass, eight longevity and five reproduction measurements, reflecting their taxonomically restricted coverage. Between-family and within-family, between-genus opportunity was available for every retained trait. Within-genus opportunity was available for 147 traits and absent for six brain traits (Supplementary Fig. 3b,d). These denominators establish which taxonomic summaries can be interpreted for each trait and prevent missing categories from being coded as observed zeros.

Supplementary Figure 3a displays the balance of Upper and Lower tail counts at each estimable level, while panel c compares the observed tail fraction with the fraction of all available pairs at that level. Their signed vertical difference is the opportunity-adjusted excess reported in Main Figure 2: positive values indicate over-representation relative to opportunity, zero proportional representation and negative values under-representation. This adjustment permits comparisons among traits with different species coverage, but it remains descriptive. A large excess does not demonstrate a departure from BM or OU, and a small excess does not demonstrate agreement with either model.

##### 3.4. Evolutionary-model selection and bootstrap stability

BM and OU fits were compared separately for every retained trait. The operational rule selected OU when AIC(BM) - AIC(OU) exceeded 2; otherwise BM was retained. Ninety-two traits selected BM and 61 selected OU (Supplementary Fig. 4a). The association between model support and matched-species coverage is displayed explicitly because sparse traits can provide limited information for distinguishing the two processes. For OU fits, the attraction parameter alpha was inspected together with relative model support; values near the numerical boundary identify OU fits that approach BM-like behaviour and should not be interpreted as strong evidence for attraction merely because an OU model was fitted (Supplementary Fig. 4b).

Model stability was measured as the fraction of 1,000 full parametric-bootstrap replicates in which refitting and reselection retained the generating model selected from the observed data. Stability ranged from 0.497 to 1.000. Eighteen traits were below 0.80, 17 ranged from 0.80 to less than 0.90, 86 ranged from 0.90 to less than 0.95 and 32 were at least 0.95 (Supplementary Fig. 4c). The horizontal lines at 0.80 and 0.95 are descriptive landmarks rather than inferential thresholds. Stability measures reproducibility of the BM/OU choice under the fitted generating process; it is not itself evidence for an unusual PSS depth distribution.

#### 3.5. Full null calibration of phylogenetic-depth profiles

The complete inferential bootstrap retained every operation that could alter the observed result. For each of 1,000 replicates per trait, phenotypes were simulated under the selected BM or OU model, both models were refitted and reselected, all pairwise PSS values were recalculated and the same Upper and Lower 1%

definitions were reapplied. The cumulative depth profile from each replicate therefore incorporates uncertainty arising from stochastic trait evolution, model fitting, model choice, reuse of species across pairs and the score-dependent selection of the empirical tails.

The atlas in Supplementary Figure 4d uses a root-to-tip coordinate from 0 (rootward and deep) to 1 (tipward and shallow). This should not be confused with the within-trait opportunity percentile used for the raw-depth summary in Main Figure 2a, where the shallowest and deepest available comparisons define the ends of a rank scale. Each atlas cell represents the observed cumulative profile minus the median full-bootstrap profile. Blue values denote deepward redistribution, red values shallowward redistribution and black marks identify depth intervals outside the trait-specific simultaneous 95% envelope.

Directional signed-area probabilities were corrected across traits using the Benjamini-Hochberg procedure. At FDR q <= 0.10, 20 Upper 1% profiles showed supported deepward redistribution and two Lower 1% profiles showed supported shallowward redistribution (Supplementary Fig. 4e). The 20 Upper-tail results comprised 11 body-mass measurements, five brain measurements, three biometabolite traits and territoriality. Related body- and brain-size definitions are correlated measurements and demonstrate robustness across alternative definitions, not independent biological replications. Simultaneous-envelope crossing provides complementary localization of the depth intervals contributing to a profile, but neither the envelope marks nor the signed-area result assign significance to individual species pairs, nodes or branches.

##### 3.6. Interpretation, reporting boundaries and data release

Together, Supplementary Figures 2-4 establish an audit trail from the source phenome to the final trait-level inference. Trait filtering defines the analysable variables, phylogenetic matching defines the available pair universe, opportunity adjustment separates representation from sampling geometry and the full refitted bootstrap tests aggregate redistribution relative to the fitted evolutionary process. The principal inferential unit is therefore the complete trait-by-tail depth profile. Empirical tail membership remains a discovery device that preserves species identities for downstream localization and genome-phenome hypothesis construction.

The figures intentionally summarize rather than print every per-trait record. The complete retention audit, family coverage matrix, exclusive opportunity counts, model diagnostics and bootstrap statistics are available in the machine-readable files used to construct the figures. During final Supplementary Information assembly, compact items needed for reading should remain conventional Supplementary Tables, whereas exhaustive trait-by-tail audits should be released as Source Data. This separation keeps the printed Supplementary Material interpretable while preserving complete reproducibility of the phenome-wide analysis.

#### Supplementary Note 4. Cross-scale localization of brain-body differentiation from mammals to Cercopithecidae

This note provides the analytical bridge between the mammalian brain-body dataset and the nested primate case study used to evaluate Phenotype Shift Scores (PSSs). The analysis proceeds across three scales. First, body mass, brain mass and allometry-adjusted brain mass are compared across mammals. Second, a mammal-wide phylogenetic allometric model defines a signed residual that is retained as a common phenotype in Primates. Third, PSS is refitted within the primate phylogeny and then within Cercopithecidae to localize high-ranking comparisons without redefining the phenotype at each taxonomic level. Supplementary Figures 5-7 document, respectively, the mammalian taxonomic distributions, diagnostic assessment of the allometric phenotype and nested localization in primates and cercopithecids.

##### 4.1. Mammalian data and construction of the three brain-body phenotypes

Species-level brain and body masses were obtained from the mammalian dataset assembled by Venditti, Baker and Barton (2024). The source study collated measurements from the literature, prioritizing brain mass over converted volume and, where possible, paired brain and body measurements from the same individuals. After the project-specific preparation, taxonomic reconciliation and exact matching to the mammalian phylogeny, 1,427 species had finite positive values for both variables. Masses were expressed in grams before transformation, and natural logarithms were used throughout.

We analysed three related continuous phenotypes: log body mass, log brain mass and allometry-adjusted brain mass. The adjusted phenotype was defined as the signed residual from a mammal-wide phylogenetic quadratic regression of log brain mass on log body mass. The reference model was fitted by maximum likelihood with Pagel's lambda using phylolm (Pagel, 1999; Ho and Ané, 2014): log brain mass = -2.35631 + 0.714793(log body mass) - 0.0110693(log body mass)$^2$. The fitted phylogenetic-signal parameter was lambda = 0.978887. Positive residuals therefore identify species with greater brain mass than predicted by the mammal-wide curve at their body mass, whereas negative residuals identify species below that expectation.

This residual is an operational, body-size-adjusted phenotype; it is not an estimate of cognitive capacity, regional neuroanatomy or evolutionary rate. It was calculated once from the mammalian reference and then transferred unchanged to nested analyses. Consequently, any primate or family-level result retains the same mammalian baseline and cannot arise from repeatedly re-centring the phenotype within progressively narrower clades.

##### 4.2. Mammalian comparison of body mass, brain mass and allometry-adjusted brain mass

Each phenotype yielded 1,017,451 unordered species pairs, of which 10,175 formed the empirical Upper 1% of its PSS distribution. These tails were used as trait-specific prioritization sets, not as collections of statistically significant pairs. Supplementary Figure 5 compares their order-level composition, representation relative to pairwise opportunity, within-order score distributions and taxonomic coverage. Across the source taxonomy, 29 order-level labels were retained, and 24 contained at least one within-order pair. The strong variation in species coverage and in the number of available within-order comparisons makes the opportunity panels essential for interpreting raw upper-tail counts.

The three upper tails were not taxonomically interchangeable. Among Upper 1% pairs involving Primates, 416 of 431 body-mass pairs (96.5%) and 519 of 521 brain-mass pairs (99.6%) were within Primates. For allometry-adjusted brain mass, 1,057 of 1,576 primate-involving pairs were within the order (67.1%); the remaining comparisons connected a primate with another mammalian order (Supplementary Fig. 5a). Comparable trait-specific shifts in within- versus between-order composition occurred in Carnivora, Rodentia, Cetacea and other well-sampled groups.

The endpoint-based opportunity analysis further separated abundance from representation. For each order, the observed number of Upper 1% endpoints was compared with all endpoints available to its member species; a within-order pair contributed two endpoints. This revealed marked differences among orders and traits even after accounting for the number of represented species (Supplementary Fig. 5b,d). The density panel shows that those differences were not reducible to a uniform shift in pair abundance, because the shapes of within-order PSS distributions also varied among traits (Supplementary Fig. 5c). These summaries locate the empirical prioritization sets; they do not by themselves imply an order-specific rate increase or a departure from the fitted BM/OU null.

Full refitted-bootstrap calibration, reported in the main Results, showed that the Upper 1% depth profiles also differed among the three phenotypes. Signed observed-minus-null areas were -0.118 for log body mass (FDR-adjusted q = 0.059), -0.094 for log brain mass (q = 0.119) and -0.159 for allometry-adjusted brain mass (q = 0.030). Negative values denote redistribution towards deeper comparisons. Because adjusted brain mass is derived from the two raw measurements, these outcomes are a structured comparison of related phenotypes rather than independent biological replications. Their contrast indicates that removing the mammal-wide body-size relationship exposes phylogenetic-depth structure not supported for raw brain mass alone.

##### 4.3. Specification and diagnostic assessment of the mammal-wide allometric phenotype

The allometric regression is a phenotype-construction model and is analytically distinct from the BM and OU models subsequently fitted to calculate PSS. Its adequacy is evaluated in six prespecified diagnostic blocks (Supplementary Fig. 6). First, the frozen quadratic lambda model is refitted and checked against the stored coefficients, lambda, residual variance and log likelihood. A 400-point prediction grid represents the fitted mean relationship and its pointwise 95% confidence interval conditional on the maximum-likelihood lambda estimate. Second, raw residuals are checked against the stored allometry-adjusted phenotype and examined alongside residuals whitened by the fitted Pagel-lambda covariance.

Third, residual magnitude and scale-location patterns are evaluated against fitted values and body mass using rank associations and robust descriptive slopes. Fourth, generalized leverage, studentized residuals and Cook distances are calculated after phylogenetic whitening, followed by exact deletion refits for *Homo sapiens* and prespecified influential observations. Fifth, linear, quadratic and cubic lambda models are compared using AICc, Akaike weights and nested likelihood-ratio contrasts, with agreement of the resulting residual tails assessed directly. Sixth, residual stability is quantified under removal of *Homo sapiens*, all Hominidae, the 15 largest Cook distances and substitution of the complete linear model. These checks use one frozen mammalian phylogeny and therefore test model-form, influence and taxonomic sensitivity rather than uncertainty among alternative mammalian trees.

The reference quadratic fit explained 82.7% of the variance in log brain mass and retained strong phylogenetic signal (Pagel's lambda = 0.979; Supplementary Fig. 6a). Its diagnostics nevertheless identified departures from a homoscedastic Gaussian error description. Phylogenetically whitened residuals had an excess kurtosis of 3.76, and their quantile plot showed heavier tails than expected under normality (Supplementary Fig. 6b). Absolute residual magnitude increased with fitted brain mass and log body mass (Spearman's rho = 0.427, P = $2.95 \times 10^{-64}$), producing the positive scale-location trend in Supplementary Fig. 6c. These patterns do not alter the definition of the signed residual, but they show that the quadratic model is an imperfect description of residual dispersion.

Influence was concentrated in a small number of morphologically extreme mammals, particularly cetaceans. *Balaena mysticetus* and *Lissodelphis borealis* had the two largest Cook distances (0.728 and 0.510, respectively); deletion of *L. borealis* changed the quadratic coefficient by 29.9% and produced the largest local change in the fitted curve. By contrast, *Homo sapiens* ranked 24th by Cook distance, and its deletion had only a modest effect on predictions (leave-one-out prediction RMSE = 0.0028; Supplementary Fig. 6d). The allometric phenotype is therefore not driven by humans or hominids, although a few large-bodied non-primate species exert appreciable leverage on the mammal-wide curve.

Model-form comparison did not identify the original quadratic specification as the best-supported relationship. The cubic Pagel-lambda model had the lowest AICc, whereas the quadratic and linear models had Delta AICc values of 31.0 and 149.5, respectively; adding the cubic term improved fit over the quadratic model (likelihood-ratio test = 33.05, 1 d.f., P = $8.99 \times 10^{-9}$; Supplementary Fig. 6e). This statistical improvement caused comparatively limited global reordering: quadratic and cubic residuals had Spearman's rho = 0.9977, with 96.5% overlap in the Upper 10% and 88.1% overlap in the Lower 10%. Nevertheless, 32 species changed residual sign and membership near empirical tail thresholds was not invariant to model form.

The prespecified sensitivity analysis supported the taxonomic stability of the focal signal while clarifying its model dependence (Supplementary Fig. 6f). Excluding *H. sapiens* or all Hominidae left primate residual ranks virtually unchanged (Spearman's rho = 0.999986 and 0.999973, respectively) and preserved both 10% tails. Removing the 15 largest Cook distances retained high rank agreement but reduced Upper 10% overlap to 90.5% in Primates and 88.9% in Cercopithecidae. Replacing the quadratic model with the poorly supported linear form caused larger tail changes, with Upper 10% overlaps of 85.7% and 88.9% in the same groups. These values refer to residual rankings and phenotype tails, not to a refitted PSS analysis.

Taken together, the diagnostics provide mixed rather than unqualified validation. They support the need for allometric adjustment and show that the nested primate result is not an artefact of *H. sapiens* or Hominidae, but they do not establish the quadratic specification as uniquely or optimally supported. The present PSS analyses retain the quadratic residual as the declared reference phenotype. Exact PSS pair identities, especially close to the 1% thresholds, should remain conditional on this choice until the score is recomputed from cubic-model residuals. More generally, diagnostic support for a signed residual would not establish that it captures all biologically relevant dimensions of encephalization.

##### 4.4. Transfer of the mammalian residual and PSS refitting in Primates

The prepared mammalian table contained 204 species classified as Primates. Of these, 129 matched exactly to tips in the fossil-calibrated S4 phylogeny distributed by Kuderna et al. (2023); the tree was pruned to those taxa while preserving its temporal branch lengths. The mammal-wide residuals were copied to the matched species without re-estimation. The retained primate dataset therefore combines a class-wide allometric reference with a primate-specific evolutionary covariance structure.

BM and single-optimum OU models were refitted to the 129 residuals, OU was selected, and PSS was calculated for all 8,256 unordered pairs. Eighty-three pairs formed each empirical 1% tail. Upper-tail comparisons were assigned once to the most recent common ancestor (MRCA) of their two endpoints, enabling the pair-ranking output to be localized on the primate tree while retaining species identities. Node fill and family-level summaries were normalized by the number of pairs available to each node or taxonomic group. These quantities describe concentration relative to sampling opportunity; they are not node- or family-specific hypothesis tests.

Supplementary Figure 7a provides the complementary localization of the 83 Lower 1% pairs using the same MRCA and opportunity conventions as the upper-tail main figure. Showing both tails is important because low-ranking pairs are not simply the inverse of high-ranking pairs: the score combines observed magnitude, model-based placement and phylogenetic distance, and the available pair geometry differs among nodes.

##### 4.5. Family localization and the within-Cercopithecidae refit

The primate Upper 1% contained 43 within-Cercopithecidae pairs, compared with 14.929 expected from the family's share of all within-family pair opportunity (Supplementary Fig. 7b). This corresponds to an observed-to-expected ratio of approximately 2.88. Other families contained between zero and five within-family Upper 1% pairs. Some small families showed larger ratios but contributed few pairs because their opportunity was limited. Cercopithecidae was therefore selected as a high-information focal domain because it combined a concentrated upper-tail signal with 55 matched species and 1,485 available within-family comparisons, not because it uniquely maximized an enrichment statistic.

Across the 30 primate genera represented by at least two species, 13 genera contributed 45 congeneric comparisons to the 83-pair Upper 1% (54.2%; Supplementary Fig. 7c). The largest contributions were 21 of 78 available comparisons in *Macaca*, seven of 28 in *Trachypithecus*, four of ten in *Papio* and two of three in *Colobus*. These counts jointly display observed localization and the sharply different number of possible comparisons among genera. They should not be interpreted as genus-specific significance tests.

We next repeated the complete model-fitting and scoring procedure within Cercopithecidae. The 55-species tree yielded 1,485 unordered pairs, OU was again selected and 15 pairs formed the empirical Upper 1%. Fourteen of those 15 comparisons were congeneric (Supplementary Fig. 7d). The highest-scoring congeneric comparison joined *Trachypithecus cristatus* and *T. auratus*; high-ranking genus-level contrasts also included *Macaca nigra* versus *M. tonkeana*, *Papio anubis* versus *P. papio*, and *Colobus polykomos* versus *C. guereza* (Supplementary Fig. 7e). The separate refit is critical: it asks which comparisons rank most highly under the evolutionary and scaling context of the focal family rather than merely subsetting the primate-wide ranking.

A fixed-tail leave-one-species-out analysis evaluated how the primate localization summaries changed when each species was removed without refitting PSS (Supplementary Fig. 7f). Across the 129 deletions, the congeneric share ranged from 50.0% to 76.3%, and the Cercopithecidae observed-to-expected ratio ranged from 2.65 to 3.99. The largest excursion occurred after removal of *Homo sapiens*, illustrating that this panel is sensitive to changes in the retained fixed tail and its total opportunity. Because neither the evolutionary model nor pair rankings were recomputed, this analysis is a deletion diagnostic for the observed catalogue, not a full robustness refit.

##### 4.6. Cross-scale interpretation and reporting boundaries

The three supplementary figures establish a continuous analytical chain. Supplementary Figure 5 shows that body mass, brain mass and their mammal-wide residual generate different taxonomic compositions and PSS distributions. Supplementary Figure 6 quantifies the diagnostic strengths and limitations of the residual used as the focal phenotype, including leverage and model-form dependence. Supplementary Figure 7 then preserves the declared quadratic phenotype while changing only the phylogenetic and scoring scope, thereby localizing candidate comparisons from Primates to Cercopithecidae and to individual genera.

This nested strategy avoids two forms of circularity. The phenotype is not recalculated to accentuate variation in the clade selected for downstream analysis, and the within-family PSS result is obtained by refitting the declared evolutionary models rather than manually selecting visually extreme species. At the same time, neither PSS nor the localization summaries identify the causal branch, establish convergent molecular evolution or test a genome-phenome mechanism. They formulate explicit candidate contrasts. Their added value is evaluated downstream by comparing PSS-informed foreground/background groups with non-PSS hypotheses, including absolute top/bottom phenotype selections, under the same genome-scale analysis framework.

The principal biological limitation is equally important. Total allometry-adjusted brain mass is a coarse integrative phenotype. It does not isolate cortical regions, neuronal composition, developmental timing or cognitive function, and its signed residual depends on the mammalian reference model and source measurements. Conclusions should therefore remain at the level of relative brain-mass differentiation until independent anatomical, ecological or molecular evidence supports a more specific interpretation.

#### Supplementary Note 5.Pooled CAAStools validation of PSS-informed genome-phenome hypotheses

This note evaluates whether Phenotype Shift Scores (PSSs) improve the formulation of genome-phenome hypotheses for a continuous trait. All strategies used the same allometry-adjusted brain-mass phenotype, the same primate phylogenetic framework and the same pooled CAAStools analysis, but differed in how foreground (FG) and background (BG) species were selected. The comparison therefore asks whether PSS-informed selection yields a more focused and functionally interpretable candidate set than phenotype-only or taxonomically defined alternatives. Supplementary Figure 8 documents strategy composition, pooled-cycle sampling and sequence-level quality control; Supplementary Figure 9 compares the resulting gene sets and functional enrichments.

##### 5.1. Definition of the four genome-phenome strategies

Four canonical strategies were retained for direct comparison. In every strategy, species with higher allometry-adjusted brain mass were assigned to FG and species with lower values to BG. Each pooled hypothesis contained four FG and four BG species. The direction of the molecular comparison was therefore held constant, while the biological rule used to construct the candidate groups varied. Exact species membership and phenotype values are reported in the source data underlying Supplementary Figure 8a.

**Primate-wide extremes.** The phenotype-only, broad-scale strategy used the upper and lower absolute tails of the allometry-adjusted brain-mass distribution across primates. The fixed discovery pools contained 13 FG and 13 BG species. Four species were drawn independently from each side, with no repeated genus within FG or within BG. This design represents the common practice of contrasting distant species selected solely because they occupy opposite ends of a continuous phenotype distribution.

**Family-level parallel pairs.** This taxonomically structured strategy defined one directional contrast per informative primate family: the species with the highest relative brain mass entered FG and the species with the lowest value entered BG. The resulting 13 paired family contrasts provided 13 FG and 13 BG species. Each cycle sampled four complete family pairs, thereby preserving a repeated within-family high-versus-low contrast while distributing the comparison across the primate tree.

**Cercopithecidae PSS-driven groups.** The PSS-informed strategy was restricted to the 55-species Cercopithecidae analysis described in Supplementary Note 4. A pair from the empirical Upper 1% of the within-family PSS distribution contributed to pool definition when at least one of its endpoints also occurred in a Lower 1% pair. This rule identifies species participating both in unusually differentiated and unusually similar comparisons, using the score to define informative positions in the phenotypic network rather than selecting absolute phenotype endpoints. Within each retained upper-tail pair, the higher-phenotype endpoint was assigned to FG and the lower endpoint to BG. Deduplication produced fixed pools of nine FG and ten BG species with no species assigned to both sides. Pair membership justified inclusion in the pools but did not constrain individual CAAStools cycles: four species were sampled independently from each side.

**Cercopithecidae-wide extremes.** The closest phenotype-only comparator applied an upper/lower 10% rule after restricting the same phenotype table to Cercopithecidae. Rounding the tail size upward yielded six FG and six BG species. Four species were sampled independently from each side without a genus restriction. This control is particularly informative because it matches the PSS strategy's taxonomic domain while replacing its pairwise, phylogenetically contextual selection rule with absolute within-family tails.

The comparison was not designed to maximize the number of significant genes or functional terms. Broad trait tails can include most of the tested background and therefore produce many sequence candidates with limited selectivity. Instead, the prespecified criterion was whether PSS generated a tractable subset that retained repeated FG/BG support and showed coherent functional structure relative to the same-family absolute-tail control.

#### 5.2. Construction and coverage of pooled sampling cycles

For every strategy, 100 unique 4-versus-4 cycles were selected without replacement from the admissible set. Species could recur across cycles but not within a cycle. A fixed seed (260821) and deterministic SHA-256 ranking made cycle selection reproducible across software environments. The cycles are alternative realizations of each biological grouping rule and should not be treated as 100 independent evolutionary replicates. Their purpose is to reduce dependence on any single foreground/background draw and to reconstruct support across the finite discovery pools.

The number of admissible cycles differed substantially among strategies (Supplementary Fig. 8c). The selected cycles covered 100 of 233,967 possible primate-wide comparisons (0.0427%), 100 of 715 family-pair combinations (14.0%), 100 of 26,460 PSS-driven Cercopithecidae comparisons (0.378%) and 100 of 225 Cercopithecidae-wide tail comparisons (44.4%). These fractions reflect the combinatorial geometry of the fixed pools and sampling restrictions; they are not measures of analytical quality.

Per-species sampling frequencies were inspected against the expectation implied by 400 FG and 400 BG selections per strategy (Supplementary Fig. 8b). In the 13-species primate-wide pools, observed frequencies ranged from 18% to 44% in FG and from 16% to 43% in BG around an expectation of 30.8%. The paired-family design was more even (23-36% on both sides). Frequencies ranged from 38% to 51% among the nine PSS FG species and from 34% to 45% among the ten PSS BG species, close to expectations of 44.4% and 40.0%, respectively. In the six-species Cercopithecidae-wide pools, the corresponding ranges were 61-71% and 59-70% around an expectation of 66.7%. No duplicated cycle or discrepancy between declared and observed pool size was detected.

#### 5.3. Pooled CAAStools execution and event aggregation

Pooled convergent amino-acid substitution analyses were performed with the pooled-discovery implementation of CAAStools (Barteri et al., 2023). A Nextflow DSL2 workflow crossed each strategy with the complete collection of orthologous protein alignments and submitted one alignment-by-strategy task to the SLURM cluster (Di Tommaso et al., 2017). Jobs used the bundled CAAStools implementation in a fixed project Conda environment distributed under supplementary/ in the phyloPSS repository, relaxed-PHYLIP alignments, one CPU, 2 GB of memory and a 30-min time request. Each alignment was evaluated under the same CAAStools settings, including all three substitution-pattern classes, a maximum gap fraction of 0.5 per position and at least three observed non-gap species on both FG and BG sides of an individual cycle.

CAAStools first applied its positional hypergeometric screen at nominal $P < 0.05$. Positive cycles were then aggregated into amino-acid-compatible events. A species was counted once per event even if it appeared in multiple positive cycles, whereas incompatible amino-acid signatures at the same alignment position were retained as separate events. Only primary-event records with positional $P < 0.05$ entered the consolidated analysis. No multiple-testing correction was applied at this discovery stage; the nominal filter defined candidate positions to be compared consistently across strategies. If duplicate primary records referred to the same gene and position, one was retained deterministically by lowest positional P value, highest balanced support and then highest total support.

FG and BG support were defined as the number of distinct discovery-pool species supporting an event on each side, rather than the number of positive sampling cycles. This distinction prevents repeated selection of the same species from inflating biological support. A position's balanced support was the smaller of its FG and BG support counts. For functional analysis, a gene qualified only when at least one retained position was supported by at least four distinct FG and four distinct BG species at that same position. The primary threshold thus requires support equivalent to a complete 4-versus-4 hypothesis while allowing that support to be assembled across pooled cycles.

The four strategies produced strongly different candidate volumes (Supplementary Fig. 8d,e). Primate-wide extremes yielded 888,208 nominally significant positions in 15,307 genes; family-level pairs yielded 1,923 positions in 1,399 genes; the PSS-driven strategy yielded 2,234 positions in 1,644 genes; and Cercopithecidae-wide extremes yielded 85,168 positions in 12,208 genes. At the primary balanced-support threshold after cluster filtering, the corresponding query sets contained 15,178, 242, 1,421 and 11,834 genes. The large difference between the two same-family strategies shows that taxonomic restriction alone does not explain the selectivity of the PSS-informed design.

#### 5.4. Dense-cluster filtering and functional comparison

Nominally significant positions were screened for dense local clusters before applying the balanced-support threshold. Filtering was performed separately within every strategy and gene. A position was discarded if it belonged to an interval containing at least three distinct significant positions across a span of at least three residues and with hit density of at least 0.70. This rule targets tightly packed signals that may be produced by local alignment or annotation artefacts without excluding an entire gene when isolated supported positions remain.

The primary filter retained 562,911 of 888,208 primate-wide positions (63.4%), 1,916 of 1,923 family-pair positions (99.6%), 2,170 of 2,234 PSS-driven positions (97.1%) and 76,096 of 85,168 Cercopithecidae-wide positions (89.3%; Supplementary Fig. 8f). Because most affected genes also contained positions outside a dense cluster, gene-level retention remained above 99.2% in all four strategies: only 52, one, six and 89 genes, respectively, lost all significant positions. The filter therefore altered local positional evidence much more than overall gene presence.

Functional enrichment was performed with the official g:Profiler g:GOSt API using human annotations and a strategy-specific custom background reconstructed from every tested alignment, including genes without a positive event (Kolberg et al., 2023). The analysis used g:Profiler version e114_eg62_p19_27110d83, accessed 24 August 2026. Sources comprised Gene Ontology Biological Process, Molecular Function and Cellular Component; KEGG; Reactome; WikiPathways; TRANSFAC; miRTarBase; Human Protein Atlas; CORUM; and Human Phenotype Ontology. Significance was assessed with the g:SCS procedure at adjusted $P < 0.05$.

Query-set size and enrichment strength revealed a clear selectivity trade-off (Supplementary Fig. 9a). Primate-wide extremes selected 15,178 of 16,130 tested genes (94.1% of the custom background) and produced 259 significant terms with a median fold enrichment of 1.00. Family-level pairs selected 242 of 16,129 genes (1.5%) and produced two terms with a median fold enrichment of 18.59. The PSS-driven strategy selected 1,421 of 16,132 genes (8.8%) and produced 14 terms with a median fold enrichment of 3.50. Cercopithecidae-wide extremes selected 11,834 of 16,127 genes (73.4%) and produced 80 terms with a median fold enrichment of 1.13. Relative to the same-family absolute-tail control, PSS therefore reduced the candidate set by more than eightfold while increasing median enrichment approximately threefold.

Across the four strategies, 355 significant source-term records were recovered (Supplementary Fig. 9b). Exact overlap analysis showed that 12 of the 14 PSS-associated terms were unique to the PSS-driven strategy and two were shared with the Cercopithecidae-wide control; none was shared with the primate-wide or family-pair results (Supplementary Fig. 9d). The PSS terms connected fatty-acid metabolism and biological oxidation with immune and inflammatory functions, including antigen binding and immunoglobulin complexes, canonical inflammasome and pyroptotic response, complement regulation and activation, NOD signalling, neutrophil extracellular-trap formation and broader immunoregulatory interactions. These categories constitute a functionally structured hypothesis set, but term enrichment does not establish that any individual amino-acid substitution caused the focal phenotype.

Gene-set overlaps further clarify the comparison (Supplementary Fig. 9c). The PSS set was not composed primarily of genes absent from broader scans: only six genes were exclusive to PSS, whereas most PSS genes also occurred in one or both broad-tail candidate sets. Its added value therefore lies in prioritizing a restricted, jointly supported subset of a much larger molecular search space and changing the functional composition of the retained evidence, rather than claiming a wholly separate catalogue of genes.

#### 5.5. Sensitivity and interpretation boundaries

Cluster-definition sensitivity supported the gene-level stability of the filtering step (Supplementary Fig. 8g). For the PSS-driven strategy, alternative density, span and minimum-position rules retained 96.9-98.1% of significant positions and 99.57-99.82% of significant genes. The corresponding position-level range was wider for the primate-wide strategy (51.5-75.6%), but its retained gene fraction remained 99.48-99.82%. Thus, exact position counts depend on how a dense interval is defined, whereas the presence of at least one surviving candidate position per gene is comparatively robust.

The balanced-support analysis identifies the primary four-versus-four threshold as an important, biologically interpretable operating point rather than an arbitrary display choice (Supplementary Fig. 9e). The PSS query contained 1,638 genes at a threshold of at least three species per side, 1,421 at four per side and 72 at five per side. Its Jaccard agreement with the primary set was 0.868 at threshold three and 0.051 at threshold five. Family-level pairs showed a similarly steep transition (1,398, 242 and 26 genes), whereas the two broad-tail strategies declined more gradually because their recurrent signals involved much larger fractions of the discovery pools. Functional results should consequently be interpreted for the declared four-versus-four definition rather than as threshold-free properties of a strategy.

Candidate volume, term count and enrichment magnitude capture different properties and should not be collapsed into a single performance score. The primate-wide strategy maximized breadth but selected nearly its full tested background; family pairs maximized enrichment strength but returned only two terms; the PSS strategy occupied an intermediate region with a tractable gene set and 14 coherent terms. This pattern supports the intended role of PSS as a hypothesis-formulation tool: it uses continuous-trait magnitude, phylogenetic distance and model-based expectation to focus a downstream genome-phenome comparison. It does not by itself demonstrate molecular convergence, identify a causal branch or validate a specific biological mechanism.

The comparison also remains conditional on the input protein alignments, species coverage, allometry-adjusted brain-mass phenotype, PSS pool definition and available human functional annotations. Pooled CAAStools support indicates that an amino-acid pattern recurs among distinct species selected by a strategy; it is not a substitute for site-level evolutionary modelling, experimental validation or independent replication. Within those boundaries, the same-family comparison shows that PSS can transform a diffuse absolute-tail scan into a substantially smaller and more interpretable set of genome-phenome hypotheses for a continuous trait.

### Supplementary Figures

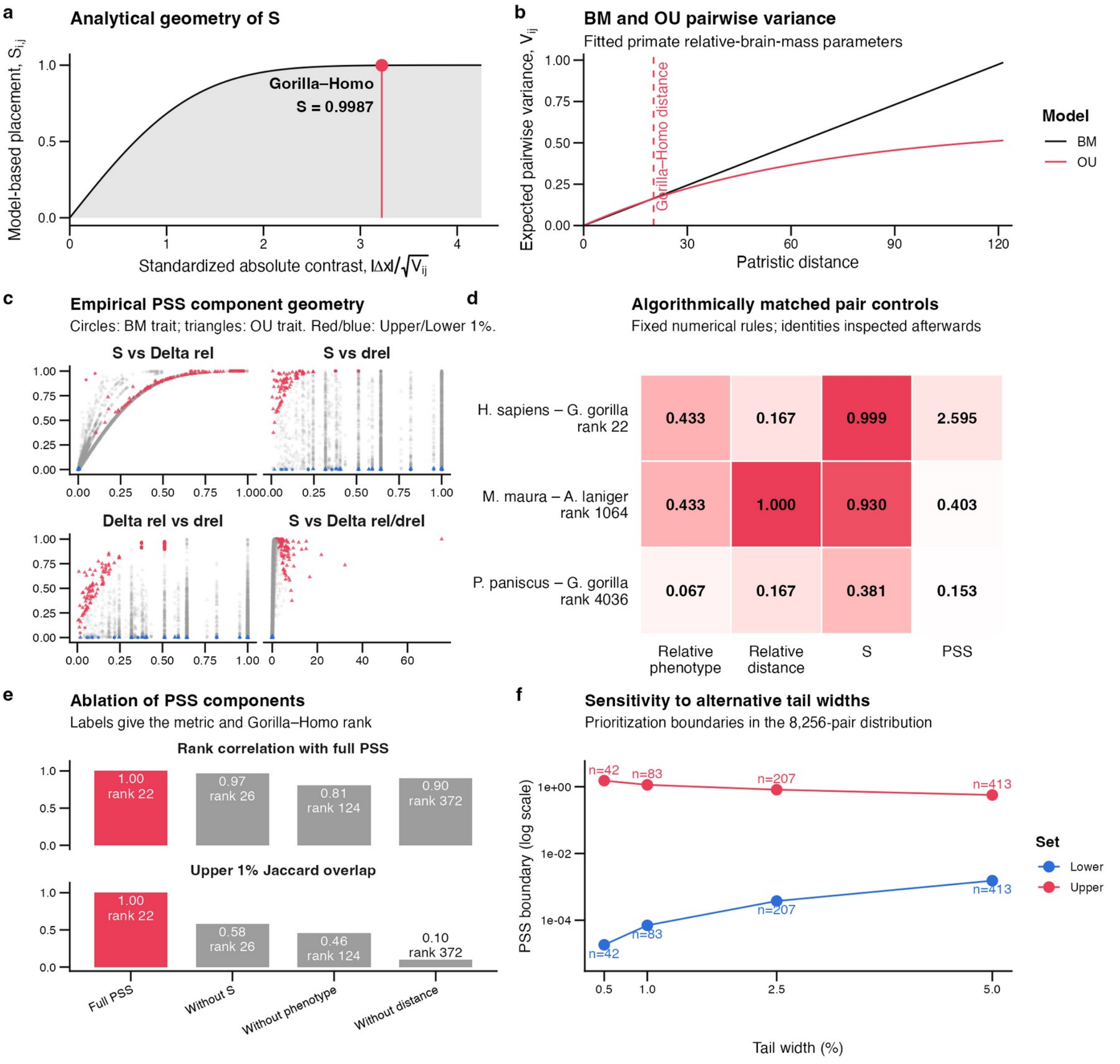


#### Supplementary Figure 1 | Component geometry and sensitivity of the phenotype shift score

The phenotype shift score (PSS) combines model-based placement, relative phenotypic difference and relative patristic distance to rank all available species pairs for a trait. **a,** Analytical relationship between the standardized absolute contrast, $z = |\Delta x|/\sqrt{V_{ij}}$, and the model-based placement score, $S_{ij} = 2\Phi(z) - 1$. The red point marks the *Homo sapiens–Gorilla gorilla* comparison for allometry-adjusted brain mass ($S$ = 0.9987). **b,** Expected pairwise contrast variance across patristic distance under Brownian motion (BM; black) and Ornstein–Uhlenbeck evolution (OU; red), using the models fitted to the 129-species primate dataset. The dashed line marks the focal pair's patristic distance. **c,** Empirical relationships among $S$, relative phenotypic difference, relative patristic distance and their ratio for male body mass, selected as BM (circles; 9,730 pairs from 140 species), and mean precipitation, selected as OU (triangles; 7,626 pairs from 124 species). Grey points denote other pairs, whereas red and blue denote the Upper 1% and Lower 1% prioritization sets, respectively. **d,** Component profiles of the focal pair and two controls selected without reference to species identity. The first control minimizes phenotypic-difference mismatch among pairs at least twice as phylogenetically distant as the focal pair; the second minimizes distance mismatch among pairs with no more than half its phenotypic difference. Cell labels give exact values; colour intensity represents the relative component value, with PSS scaled to the dataset maximum. Ranks refer to all 8,256 primate pairs. **e,** PSS component ablation. Bars report the Spearman rank correlation with complete PSS and the Jaccard overlap between Upper 1% sets after omitting $S$, phenotypic magnitude or the explicit distance denominator. Labels also give the focal-pair rank. **f,** PSS boundaries for Upper (red) and Lower (blue) prioritization sets defined at 0.5%, 1%, 2.5% and 5% of the 8,256-pair distribution. The vertical axis is logarithmic and labels give the number of selected pairs in each tail. Tail membership denotes relative prioritization, not statistical significance.

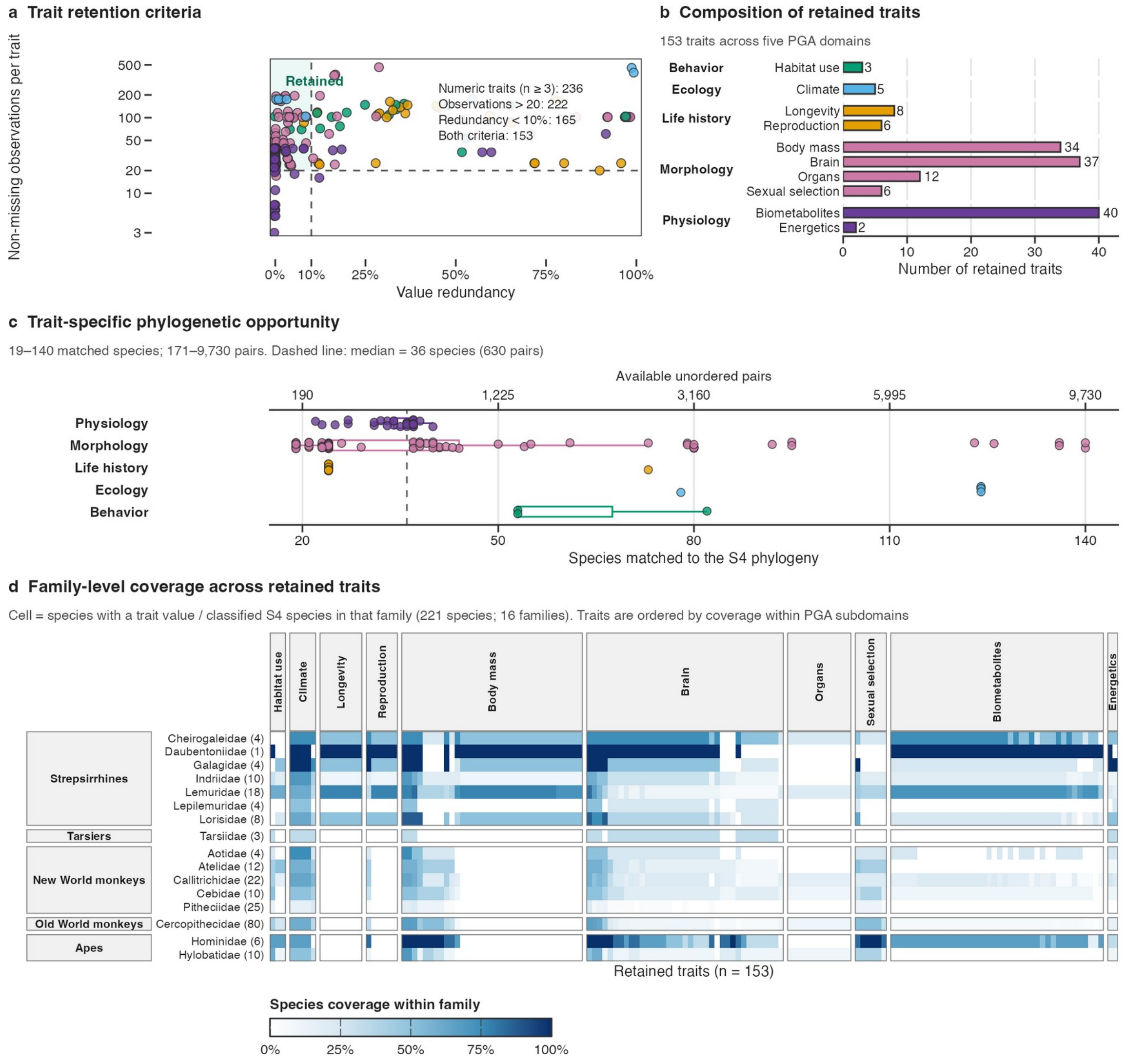


#### Supplementary Figure 2 | Trait filtering and phylogenetic coverage of the primate phenome

This figure documents the construction and effective sampling geometry of the 153-trait phenome-wide PSS analysis. **a,** Independent effects of the predefined retention criteria across 236 numeric PGA. Redundancy equals one minus the proportion of unique numeric values. Dashed lines mark the strict thresholds of more than 20 observations and less than 10% redundancy; the shaded quadrant contains retained traits. Of 236 candidates, 222 passed the observation criterion, 165 passed the redundancy criterion and 153 passed both. **b,** Retained traits by PGA domain and subdomain. Bar colours identify domains and labels give subdomain totals. The retained set comprised 3 Behavior, 5 Ecology, 14 Life history, 89 Morphology and 42 Physiology traits. **c,** Species with trait values matched to the S4 primate phylogeny. Points represent traits and colours identify domains. Box centres denote medians, boxes the interquartile range and whiskers values within 1.5 times the interquartile range. The upper axis gives available unordered species pairs, $n(n-1)/2$. Coverage ranged from 19 to 140 species and from 171 to 9,730 pairs; the dashed line marks the median of 36 species (630 pairs). **d,** Family-level coverage. Cells give the proportion of classified S4 species in each family contributing a trait value; darker blue indicates greater coverage. Rows show 16 families grouped into five major primate groups, with classified species totals in parentheses. Columns show all 153 traits, grouped by PGA subdomain and ordered by matched-species coverage. Panel **a** uses observations in the original dataset, whereas **c,d** show the phylogenetically matched species that determine pairwise opportunity in each PSS analysis.

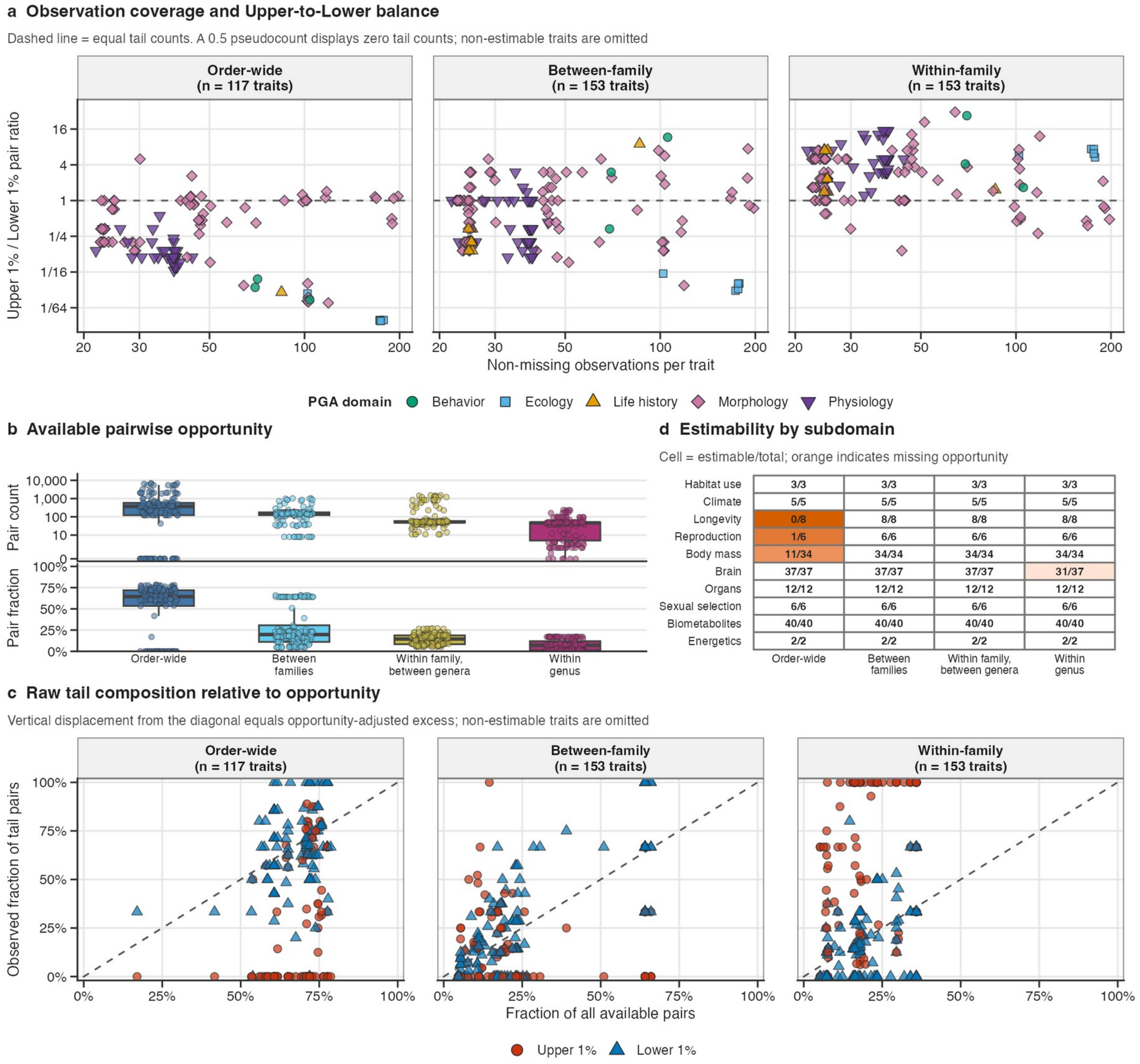


### Supplementary Figure 3 | Coverage and taxonomic opportunity across primate traits

This figure evaluates how trait coverage determines the taxonomic comparisons available to the Upper 1% and Lower 1% PSS prioritization sets. Trait data and domain assignments derive from the Primate Genome Atlas phenotype archive (PGA; Valenzuela et al., 2025). **a,** Non-missing observations versus the Upper-to-Lower pair-count ratio at the order-wide, between-family and within-family levels. Points represent traits; colours and symbols identify PGA domains. Observation counts are logarithmically scaled and ratios use a symmetric log2 scale. The dashed line denotes equal tail counts. A 0.5 pseudocount displays levels absent from one tail; traits lacking any available pair at that level are omitted. **b,** Absolute pair counts (upper plot) and fractions of all pairs (lower plot) in four exclusive categories: different major primate groups (order-wide), different families within a group, different genera within a family and species within a genus. Box centres denote medians, boxes the interquartile range and whiskers values within 1.5 times that range. Zeros are placed below one on the logarithmic count axis. **c,** Raw tail composition relative to opportunity. Points represent trait–tail combinations; red circles and blue triangles denote the Upper 1% and Lower 1%. Axes give the available and observed pair fractions, respectively. The dashed diagonal denotes equality; signed vertical displacement is the opportunity-adjusted excess used in Main Figure 2. Within-family opportunity combines the two shallowest categories in **b**. Non-estimable combinations are omitted. **d,** Estimability by PGA subdomain. Cells give estimable traits divided by the subdomain total; orange indicates incomplete opportunity. Order-wide comparisons were estimable for 117 of 153 traits, between-family and within-family, between-genus comparisons for all traits, and within-genus comparisons for 147 traits. Tail membership denotes relative prioritization, not statistical significance.

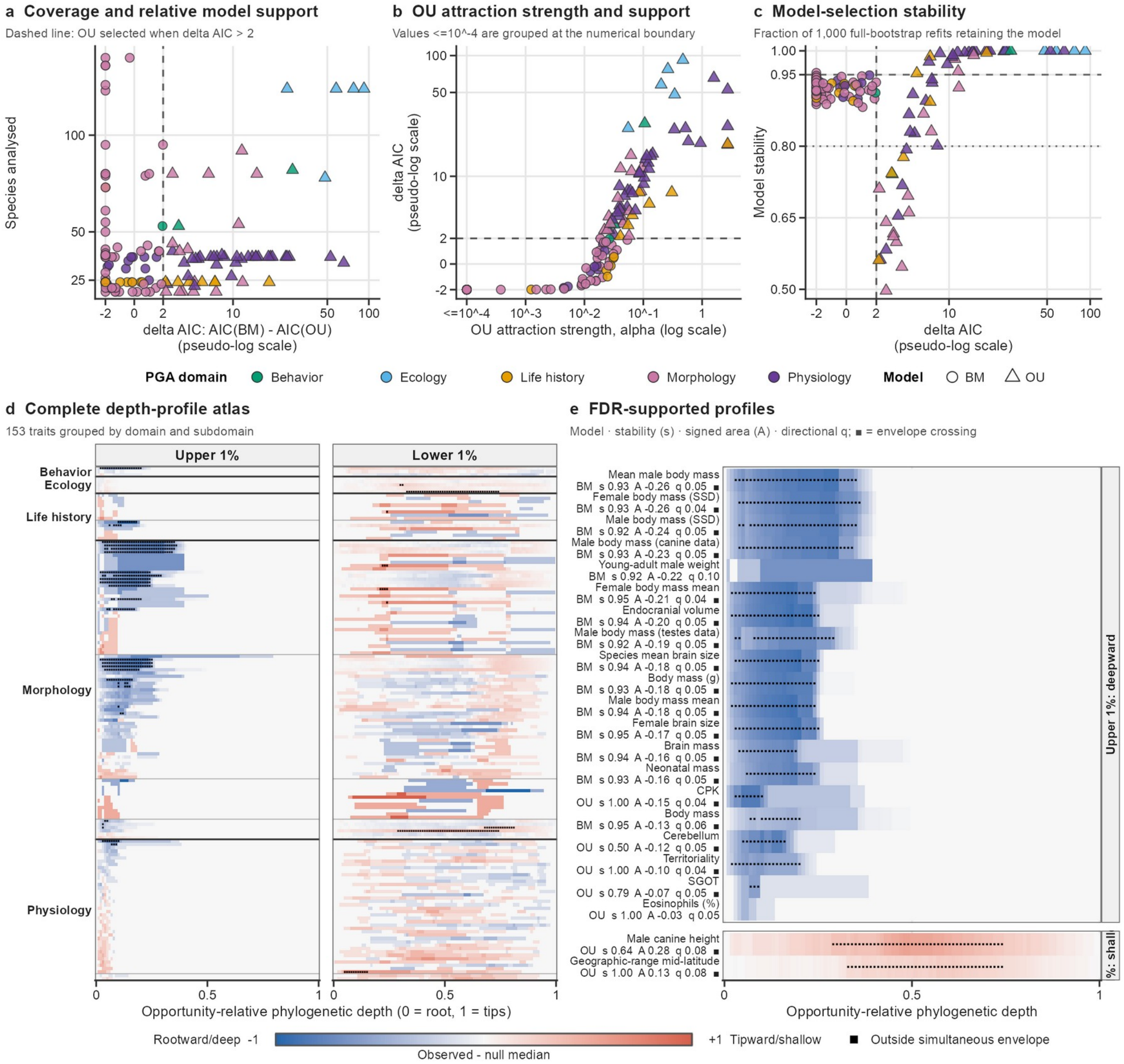


Supplementary

#### Supplementary Figure 4 | Model support, bootstrap stability and phylogenetic depth profiles across primate traits

This figure documents model selection and the full null calibration underlying the phylogenetic-depth analysis of 153 traits from thePrimate Genome Atlas phenotype archive (PGA; Valenzuela et al., 2025). **a,** Trait coverage versus relative support for Brownian motion (BM) and Ornstein–Uhlenbeck (OU) models. Each point is a trait; colour identifies the PGA domain and shape the selected model. Positive ΔAIC values favour OU, and the dashed line marks the operational OU-selection threshold, ΔAIC > 2. The ΔAIC axis uses a pseudo-logarithmic transformation. **b,** OU attraction strength, α, versus ΔAIC. The α axis is logarithmic; estimates $\leq 10^{-4}$ are grouped at the numerical boundary. The dashed horizontal line marks ΔAIC = 2. **c,** Model-selection stability, defined as the fraction of 1,000 full parametric-bootstrap replicates in which refitting retained the generating model selected from the observed trait. Horizontal lines at 0.80 and 0.95 are descriptive reference levels; the vertical dashed line marks ΔAIC = 2. **d,** Complete Upper 1% and Lower 1% depth-profile atlas. Rows represent traits grouped by PGA domain and subdomain; columns span opportunity-relative phylogenetic depth from the root (0) to the tips (1). Colour is the observed cumulative profile minus the median profile from the full refitted bootstrap. Blue denotes redistribution towards deeper comparisons and red redistribution towards shallower comparisons. Black marks identify depth intervals outside the trait-specific simultaneous 95% bootstrap envelope. **e,** Enlarged profiles passing the directional signed-area test at FDR q ≤ 0.10: 20 deepward Upper 1% profiles and two shallowward Lower 1% profiles. Labels report the generating model, model stability (*s*), signed area (*A*) and directional q value; squares mark simultaneous-envelope crossing. Related body- and brain-size definitions are correlated measurements and should not be interpreted as independent biological replications. Upper 1% and Lower 1% membership denotes relative prioritization, not statistical significance.

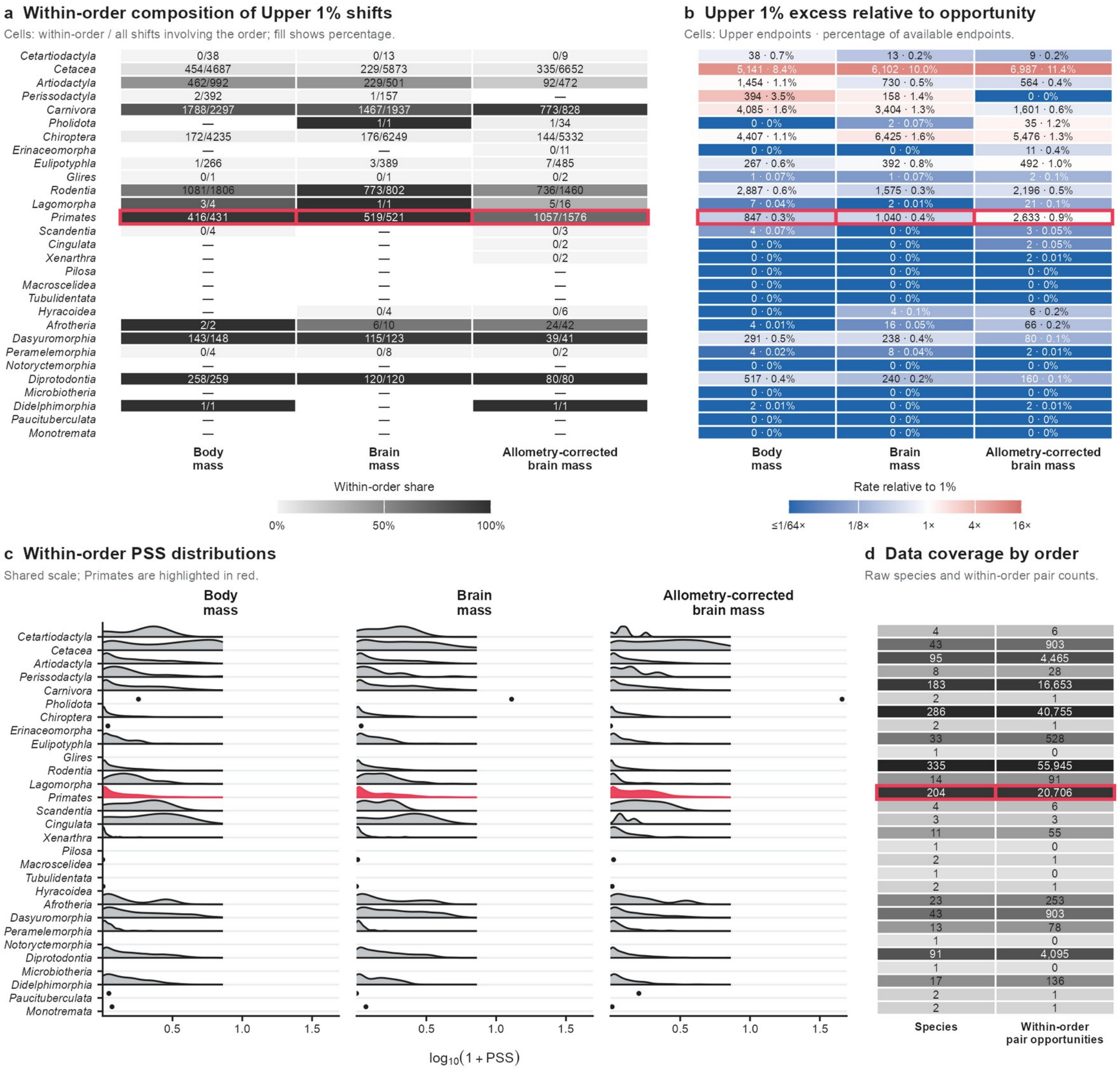


#### Supplementary Figure 5 | Order-level structure of extreme mammalian phenotype shifts

Phenotype shift scores (PSSs) were calculated for log body mass, log brain mass and allometry-adjusted brain mass across 1,427 mammals (Venditti et al., 2024), yielding 1,017,451 unordered species pairs and 10,175 Upper 1% pairs per trait. **a,** Composition of the Upper 1% for all 29 order-level labels retained from the source taxonomy. Cell entries give within-order Upper 1% pairs divided by all Upper 1% pairs involving that order; fill gives the corresponding percentage. Dashes indicate that no Upper 1% pair involved the order. **b,** Representation relative to pairwise opportunity. Entries give the number of Upper 1% endpoints and their percentage among all endpoints available to that order, calculated as the number of annotated species in the order multiplied by 1,426 possible partners. A within-order pair contributes two endpoints. Colour gives the rate relative to the 1% dataset-wide expectation; blue and red indicate depletion and enrichment, respectively. **c,** Within-order PSS distributions on a common log10(1 + PSS) scale. Each density is normalized to its own maximum and therefore compares distributional shape rather than pair abundance; isolated points denote order–trait combinations with fewer than three within-order pairs. **d,** Species coverage and the resulting number of within-order pair opportunities, $n(n - 1)/2$. Shading is scaled independently within each column and entries report raw counts. Primates are outlined or coloured in red throughout. Upper 1% membership denotes relative prioritization within each trait-specific PSS catalogue, not statistical significance. Exact numerators and denominators are provided in the accompanying order-level audit table.

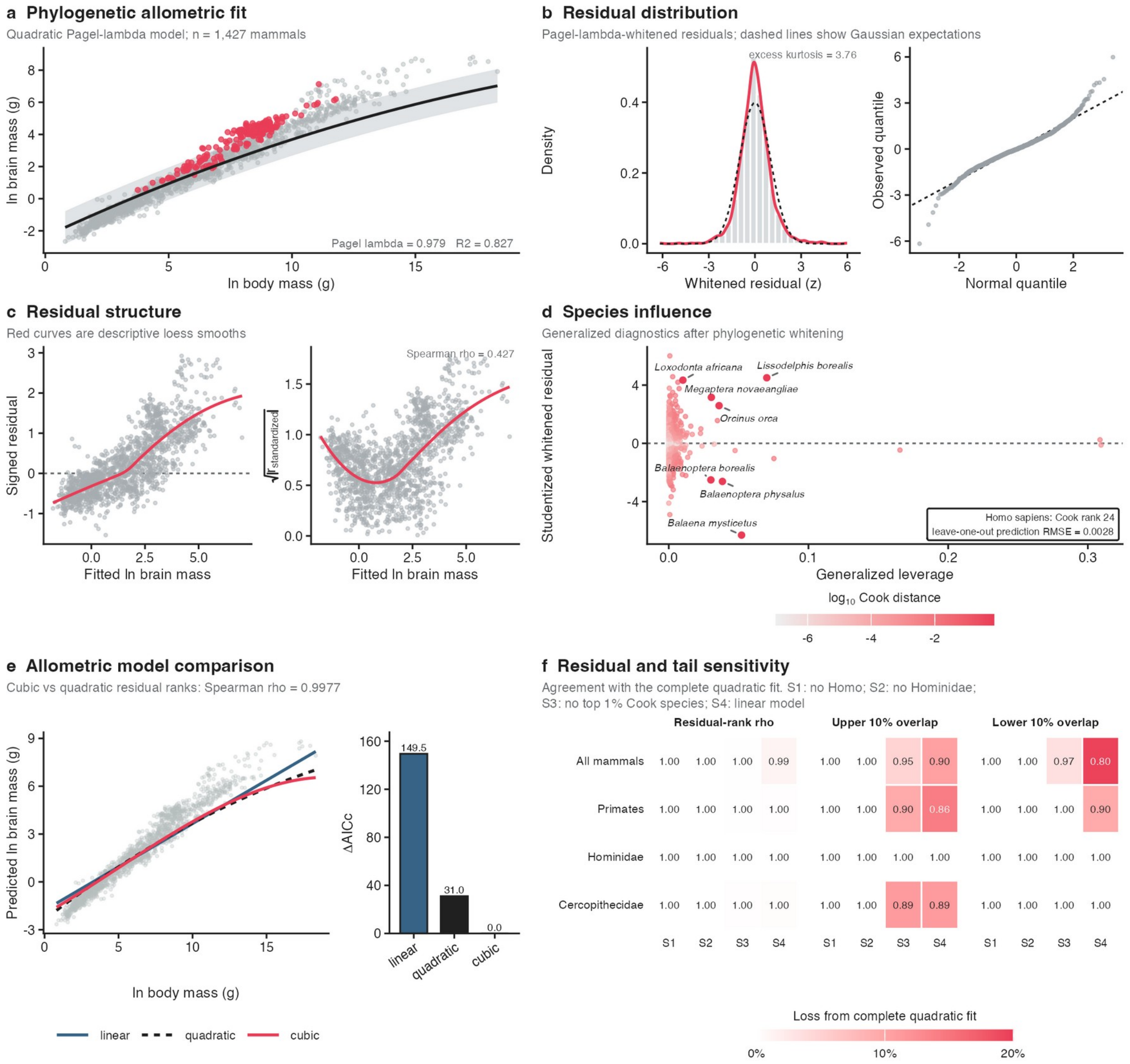


#### Supplementary Figure 6 | Diagnostics and sensitivity of the mammalian brain–body allometric model

The allometry-adjusted phenotype was derived from a phylogenetic model of log brain mass across 1,427 mammals. **a,** Observed log brain and body masses, the original quadratic Pagel-λ fit and its pointwise 95% confidence interval for the conditional mean; primates are highlighted in red. **b,** Distribution and normal quantile plot of centred, scaled residuals after phylogenetic whitening. The dashed curves or lines show Gaussian expectations. **c,** Signed residuals and the square root of the absolute standardized residual against fitted brain mass. Red loess curves are descriptive; the positive association between absolute residual magnitude and fitted mass indicates increasing dispersion. **d,** Generalized leverage and studentized residuals after phylogenetic whitening, coloured by Cook distance. The seven largest Cook distances are labelled; the inset reports the modest effect of deleting *Homo sapiens*. **e,** Prespecified linear, quadratic and cubic Pagel-λ curves and their ΔAICc values. The cubic model received the strongest support, although cubic and quadratic residual ranks remained highly concordant. **f,** Agreement with the complete quadratic fit after excluding *H. sapiens*, excluding Hominidae, excluding the 15 species with the largest Cook distances, or using a linear model. Cells report Spearman residual-rank correlations and Upper or Lower 10% overlap; colour denotes loss of agreement. Taxonomic exclusions left primate and Cercopithecidae residual ranks and tails nearly unchanged, whereas removal of influential species and linear model form had larger effects. These diagnostics concern construction of the allometry-adjusted phenotype and are distinct from the BM/OU models and bootstrap used subsequently for PSS inference. All analyses use one frozen mammalian phylogeny; alternative-tree sensitivity is not assessed here.

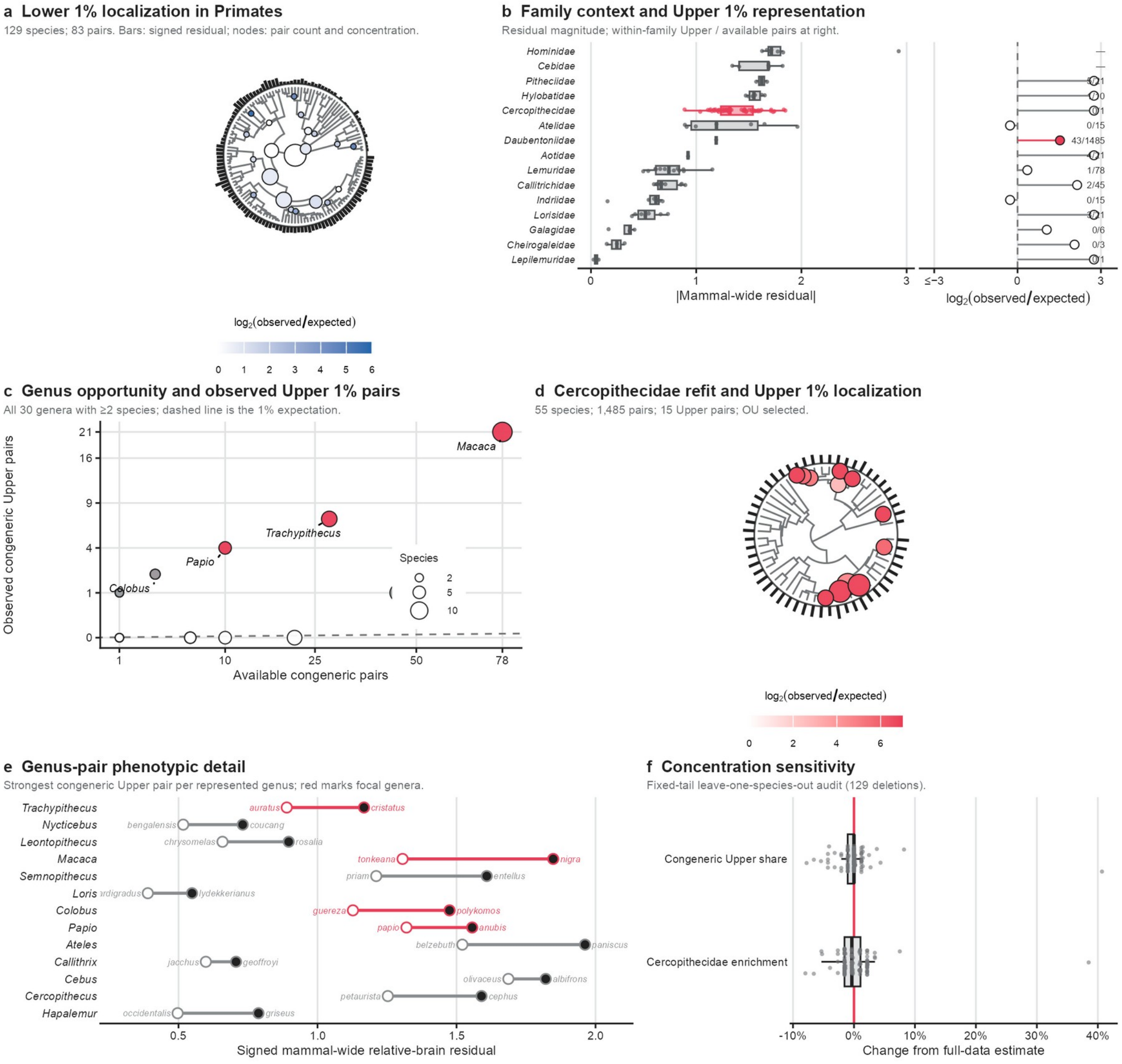


#### Supplementary Figure 7 | Nested localization of allometry-adjusted brain-mass shifts from Primates to Cercopithecidae

The mammal-wide allometry-adjusted brain-mass residual (Venditti et al., 2024) was transferred unchanged to the 129-species Kuderna S4 primate phylogeny before model refitting and PSS calculation. **a,** Localization of the 83 Lower 1% pairs. External bars give the signed residual. Each pair is assigned once to its most recent common ancestor (MRCA); node size gives pair count and fill gives log2 observed/expected concentration based on nodal pair opportunity. **b,** Family context. Left, absolute residual distributions across 15 families. Right, opportunity-adjusted within-family Upper 1% representation; labels give observed/available pairs and points give log2 observed/expected counts. Cercopithecidae, in red, contributed 43 pairs versus 14.93 expected from 1,485 available comparisons. **c,** Genus-level opportunity among all 30 genera represented by at least two species. Axes give $n(n-1)/2$ possible congeneric pairs and observed congeneric pairs in the 83-pair Upper 1%; the dashed line is the 1% expectation. Thirteen genera contributed 45 congeneric Upper pairs. **d,** PSS refit within Cercopithecidae. The 55-species analysis contained 1,485 pairs, selected an Ornstein–Uhlenbeck model and yielded 15 Upper 1% pairs, 14 of which were congeneric. Bars retain the mammal-wide residual; nodes summarize the restricted PSS refit using the encoding in **a**. **e,** Phenotypic positions of the highest-PSS congeneric pair in each represented genus. Open and filled endpoints denote lower and higher residuals; red highlights *Macaca*, *Papio*, *Trachypithecus* and *Colobus*. **f,** Fixed-tail leave-one-species-out sensitivity analysis. Each point removes one species and recalculates the congeneric Upper-pair share and Cercopithecidae enrichment from the remaining observed catalogue, relative to the full-data estimate. PSS was not refitted in this panel. Upper and Lower 1% membership denotes relative prioritization; nodal and taxonomic concentrations are descriptive rather than clade-specific significance tests.

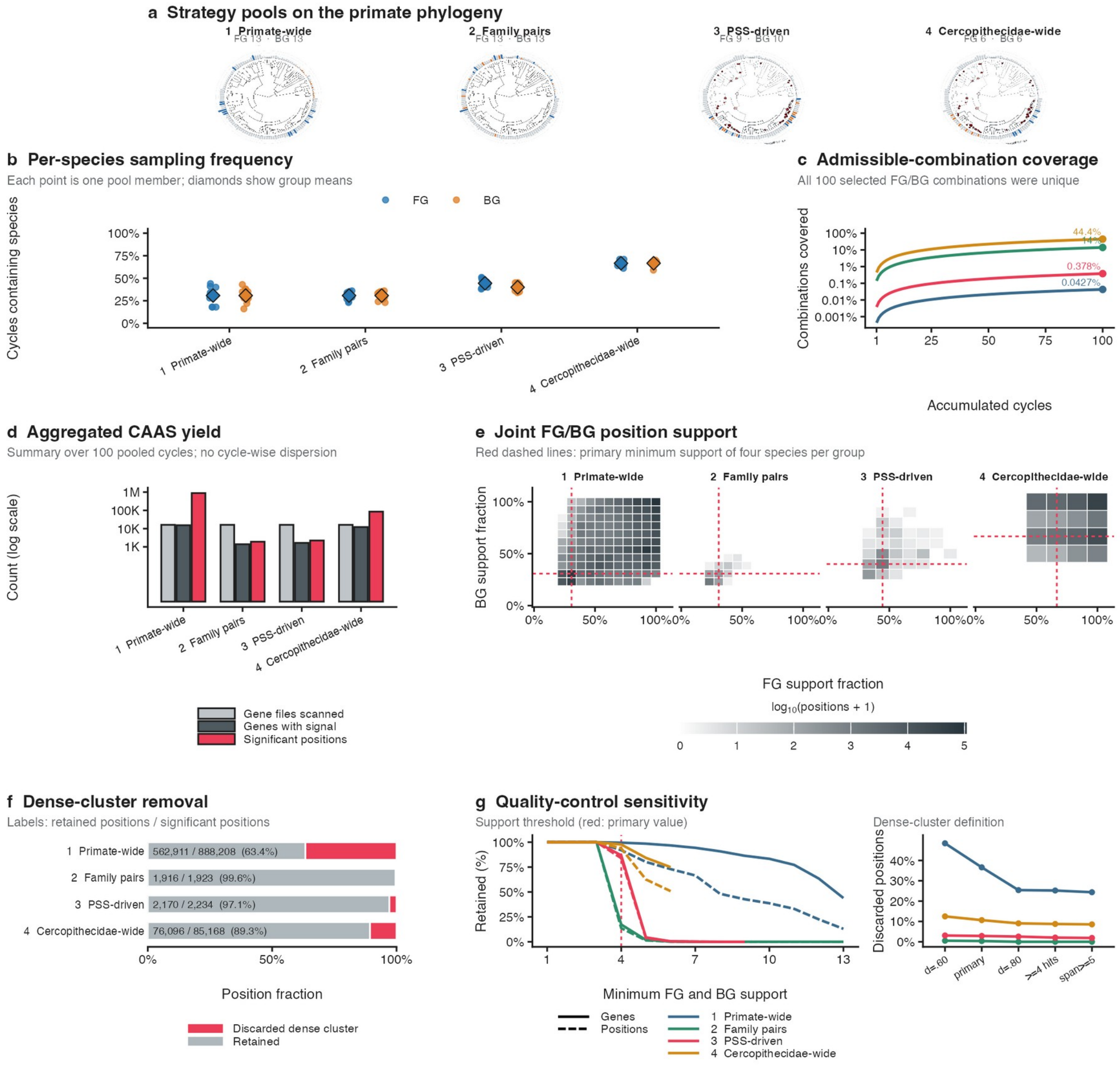


#### Supplementary Figure 8 | Sampling stability and quality control of the pooled CAAS comparison

Four foreground/background (FG/BG) strategies for relative brain mass were evaluated using 100 unique pooled CAAStools cycles, each containing four FG and four BG species. **a,** Species pools mapped onto the common 129-species primate phylogeny. External bars show relative brain mass; blue and orange identify FG and BG pool members, respectively. In the two Cercopithecidae-focused designs, dark branches delimit the family and internal circles summarize the number and opportunity-adjusted enrichment of primate-wide Upper 1% PSS shifts. **b,** Frequency with which each pool member was selected across the 100 cycles. Each point represents one species and diamonds give the mean expected from four selections per group per cycle. **c,** Cumulative coverage of admissible FG/BG combinations. All selected cycles were unique; endpoint labels give 100 divided by the strategy-specific number of possible combinations. **d,** Aggregate CAAS yield across the 100 pooled cycles: gene-level files examined, genes containing a nominally significant position and unique significant positions. Cycle-wise dispersion is deliberately not evaluated. **e,** Complete joint distributions of positional FG and BG support, normalized by the available discovery species in each strategy. Fill gives the number of positions and red dashed lines mark the primary balanced-support threshold of at least four FG and four BG species at the same position. **f,** Position retention after removal of dense local clusters, defined in the primary analysis as at least three positions spanning at least three residues with density ≥0.70. **g,** Sensitivity to the balanced-support threshold and to alternative cluster definitions. Curves show retained genes or positions relative to support threshold one; cluster-definition curves report the percentage of positions discarded. Pooled cycles are alternative samples from finite species pools and are not treated as independent biological replicates.

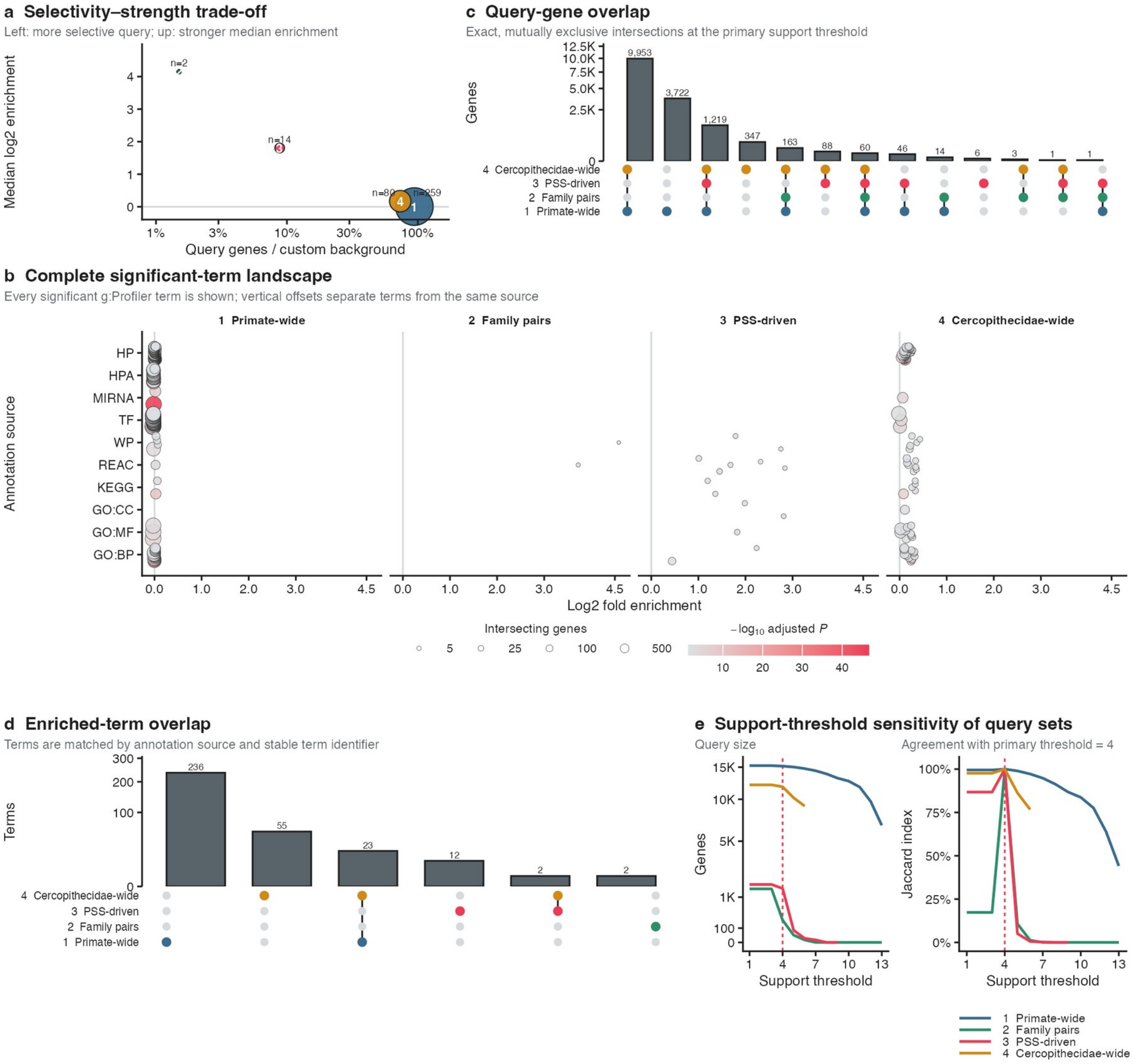


#### Supplementary Figure 9 | Complete functional comparison across pooled CAAS strategies

Functional results were compared across four foreground/background strategies using the retained genes supported by at least four foreground and four background discovery species and each strategy's tested-gene background. **a,** Trade-off between query selectivity and enrichment strength. The horizontal axis gives query genes as a percentage of the custom background, the vertical axis gives median log2 fold enrichment across significant terms and point area gives the number of significant terms. **b,** Complete landscape of all 355 significant g:Profiler results. Each point is one source–term combination; horizontal position gives fold enrichment, vertical position identifies the annotation source, colour gives adjusted significance and point area gives the number of intersecting query genes. **c,** Exact, mutually exclusive overlaps among the four primary query-gene sets. Bars give intersection size and the dot matrix identifies contributing strategies. **d,** Exact overlaps among significant terms, matched by annotation source and stable term identifier. **e,** Sensitivity of query size and composition to the balanced foreground/background support threshold. The left plot gives retained query genes and the right plot gives Jaccard agreement with the primary threshold of four; red dashed lines mark that threshold. Functional enrichment was run with g:Profiler (version e114_eg62_p19_27110d83; accessed 24 August 2026) for *Homo sapiens*, custom tested-gene backgrounds, g:SCS correction and adjusted $P < 0.05$. The archive contains no matched/randomized enrichment or alternative-background reruns; these comparisons are not inferred. Panel **e** concerns query-set sensitivity and does not represent enrichment reruns at alternative thresholds.

### Funding and acknowledgments
